# Ligand Modulation of the Conformational Dynamics of the A_2A_ Adenosine Receptor Revealed by Single-Molecule Fluorescence

**DOI:** 10.1101/2020.09.20.305425

**Authors:** Dennis D. Fernandes, Chris Neale, Gregory-Neal W. Gomes, Yuchong Li, Aimen Malik, Aditya Pandey, Alexander Orazietti, Xudong Wang, Libin Ye, R. Scott Prosser, Claudiu C. Gradinaru

## Abstract

G protein-coupled receptors (GPCRs) are the largest class of transmembrane proteins, making them an important target for therapeutics. Activation of these receptors is modulated by orthosteric ligands, which stabilize one or several states within a complex conformational ensemble. The intra-and inter-state dynamics, however, is not well documented. Here, we used single-molecule fluorescence to measure ligand-modulated conformational dynamics of the adenosine A_2A_ Receptor (A_2A_R) on nanosecond to millisecond timescales. Experiments were performed on detergent-purified A_2_R in either the ligand-free (apo) state, or when bound to an inverse, partial or full agonist ligand. Single-molecule Förster resonance energy transfer (smFRET) was performed on detergent-solubilized A_2A_R to resolve active and inactive states via the separation between transmembrane (TM) helices 4 and 6. The ligand-dependent changes of the smFRET distributions are consistent with conformational selection and with inter-state exchange lifetimes ≥ 3 ms. Local conformational dynamics around residue 229 on TM6 was measured using Fluorescence Correlation Spectroscopy (FCS), which captures dynamic quenching due to photoinduced electron transfer (PET) between a covalently-attached dye and proximal aromatic residues. Global analysis of PET-FCS data revealed *fast* (150-350 ns), *intermediate* (50-60 μs) and *slow* (200-300 μs) conformational dynamics in A_2A_R, with lifetimes and amplitudes modulated by ligands and a G-protein mimetic (mini-G_s_). Most notably, the agonist binding and the coupling to mini-G_s_ accelerates and increases the relative contribution of the sub-microsecond phase. Molecular dynamics simulations identified three tyrosine residues (Y112, Y288, and Y290) as being responsible for the dynamic quenching observed by PET-FCS and revealed associated helical motions around residue 229 on TM6. This study provides a quantitative description of conformational dynamics in A_2A_R and supports the idea that ligands bias not only GPCR conformations but also the dynamics within and between distinct conformational states of the receptor.

## INTRODUCTION

G-protein-coupled receptors (GPCRs), also referred to as seven-transmembrane (7TM) helix domain receptors, are the largest protein family in humans and are responsible for a wide range of signaling processes such as sight, smell, inflammation, cell homeostasis, and neurotransmitter and hormone mediated responses ^1^. In the human genome, GPCRs account for over 800 receptors, constituting nearly 4% of encoded proteins, and have important implications in the treatment of a wide range of diseases ^2^. In particular, the adenosine A_2A_ receptor (A_2A_R) is a class A GPCR that is important for treatment of sickle cell disease, cancer, inflammation, ischemia, neuronal disorders and various infectious diseases ^3^. Deciphering the mechanism by which GPCRs are activated and the role of ligands in stabilizing specific conformational states is essential for rational drug design.

The promiscuous interactome of most GPCRs, in addition to their reputation for exhibiting a range of efficacies as a function of ligand, implies a complex conformational landscape as revealed by a majority of recent spectroscopic studies ^4-7^. Studies of A_2A_R, rhodopsin, and a number of class A receptors have further identified a representative conformational ensemble in which ligands or interacting partners serve to stabilize specific functional states required for inactivation, G protein binding, and activation ^8-9^. Thus, ligands bias the ensemble in a manner consistent with conformational selection and facilitate allosteric transitions between these states to achieve intracellular signaling via G protein or arrestin-mediated response. ^7, 10-11^

X-ray crystallography provides snapshots of GPCRs in active or inactive states through the use of thermostabilizing mutations, fusion-proteins, or high-affinity ligands ^12^. Several crystal structures of A_2A_R have been resolved in various ligand-bound states, which show pronounced translation of the transmembrane 3 (TM3), inward shifts of TM7 and significant rotation of TM6 towards TM5 ^12-15^. Similar ligand-induced TM movements have been reported across several other class A GPCRs, including light-activatable rhodopsin GPCRs ^9 8, 16^ and the β_2_-adrenergic receptor (β_2_AR) ^17^. While X-ray crystallography does not fully capture the dynamic nature of these states, it provides a framework of key intermediates that can be used to interpret the results of techniques capable of measuring dynamics.

In recent years, several spectroscopic methods have been used to document the dynamic behavior of GPCRs, including electron paramagnetic resonance (EPR) ^11^ and nuclear magnetic resonance (NMR)^6^. Advancements in pulsed-EPR techniques, such as double electron-electron resonance (DEER), provide an opportunity to refine GPCR structures obtained by X-ray crystallography or, more recently, cryo electron microscopy ^18^, while simultaneously identifying distributions of conformations constituting the ensemble ^19-20^. Moreover, this can be repeated under various conditions (*i*.*e*., pH, ligand, detergent, liposomes, nanodiscs etc.)^21^. At the same time, EPR measurements provide a measure of dynamics of specific states, particularly on nanosecond and microsecond timescales. A combination of DEER and fluorine NMR (^19^F-NMR) spectroscopy recently highlighted the key role of orthosteric ligands in altering the free energy landscape of A_2A_R, suggesting unique activation pathways for partial and full agonists ^4 5^.

^19^F-NMR experiments using a label at the site V229C on TM6 suggest that ligand-free (apo) A_2A_R adopts two inactive states which undergo sub-millisecond exchange, resulting in a single (exchange-broadened) inactive state resonance (*i*.*e*. S_1-2_), in addition to two active-like states (S_3_ and S_3’_) which undergo slower (>5 ms timescale) exchange with the inactive state ^4^. The inverse agonist ZM241385 increases the population of the inactive states S_1-2_, the full agonist NECA preferentially stabilizes the S_3’_ activation intermediate, while the partial agonist LUF5834 increases the population of the S_3_ activation intermediate.

The binding of a peptide-mimetic of the C-terminal domain of the α subunit of G protein (Gα_s_ 374 – 394) further stabilizes the S_3’_ state in presence of NECA, while no significant changes were observed for the LUF5834-saturated receptor ^4^. The on-off flickering of the ionic lock between R102 (TM3) and E228 (TM6), which is thought to cause rapid transitions between the S_1_ and S_2_ state, was found to be suppressed in the agonist-bound state ^4, 13, 15^. A_2A_R conformational dynamics on millisecond timescales was indirectly inferred from the line-broadening of the ^19^F-NMR spectra and from Carr-Purcell-Meiboom-Gill (CPMG) relaxation dispersion measurements ^4, 22^, as well as by 2D exchange spectroscopy (EXSY) ^23^. However, a direct time-domain measurement of exchange rates between and within each of identified A_2A_R states is still lacking.

More recently, single-molecule fluorescence (SMF) techniques have been applied to GPCRs to measure their conformational heterogeneity and dynamics ^24-25^. Single-molecule Förster Resonance Energy Transfer (smFRET) is an effective tool for measuring distance distributions/changes between fluorescent dyes attached to specific sites in GPCRs within the framework of conformational heterogeneity ^26 27 28^. An smFRET study by Blanchard and coworkers on detergent solubilized β_2_AR showed evidence for ∼14 Å outward movement of TM6 and for distance fluctuations occurring on a 1-10 ms timescale ^26^. In addition, the FRET distributions were modulated by binding of orthosteric ligands and by the coupling to the G_s_ heterotrimer. Similarly, smFRET on the class C metamorphic glutamate receptor (mGluR) ^27^ revealed both millisecond (> 4 ms) and sub-millisecond (∼50-100 μs) conformational fluctuations ^28^.

Ligand-free A_2A_R has been shown to be inherently dynamic, whereby the outward motion of TM6 is important for the binding of G protein ^4^. However, the rates of these internal motions have not been directly measured. By avoiding ensemble averaging, SMF techniques have the unique ability to resolve distributions of GPCR conformers and their dynamics on a broad range of timescales^25^. Here, we used SMF methodologies that are sensitive to the movement of TM6 to measure intra-and inter-state exchange dynamics of different A_2A_R conformers. smFRET was used to measure the distribution of separation distances between TM4 and TM6 in detergent-purified A_2A_R samples labelled with a donor-acceptor dye pair at residues T119 and Q226, respectively. The smFRET distributions exhibit ligand-dependent shifts of the mean transfer efficiency and significant changes in their widths, which are interpreted by invoking three distinct conformational states of the receptor.

In addition, we labelled TM6 at residue 229 with a probe (BODIPY-FL) that is sensitive to quenching by aromatic amino acids at short-range distance (5-10 Å) ^29-30^. This process is known as Photoinduced Electron Transfer (PET) and the resulting fluorescence intensity fluctuations report on local protein dynamics around the labelling site ^31^. Fluorescence Correlation Spectroscopy (FCS) is widely used to analyze fluorescence intensity fluctuations on nanosecond-to-microsecond timescales^32^, including PET quenching processes (PET-FCS) ^30^. PET-FCS was shown to capture the kinetics of conformational dynamics in proteins, such as ultrafast folding of small water-soluble proteins ^33^ and movements of helical TM bundles in proton channels ^34^.

In A_2A_R, our PET-FCS analysis resolved three lifetimes of protein-induced quenching processes in the proximity of residue 229 on TM6: *fast* (150-350 ns), *intermediate* (50-60 μs), and *slow* (200-300 μs). The lifetimes and amplitudes of the three components are modulated by the binding of orthosteric ligands and coupling to mini-G protein (mini-G_s_). A concomitant increase in amplitude and decrease in lifetime of the fast component is observed upon activation by agonist and mini-G, while the other two components show a moderate decrease in amplitude and lifetime.

Molecular dynamics (MD) simulations of the dye-labelled receptor identified three tyrosine residues as the being responsible for the (sub)microsecond PET quenching observed by FCS: Y112 on TM3, and Y288 and Y290 on TM7. MD simulations also revealed novel helix-to-loop dynamics near the intracellular-end of TM3, which decreased significantly when the A_2A_R is prearranged to bind mini-G_s_. This work shows that GPCR ligands modulate not only state populations and associated exchange dynamics of the ensemble, but also internal dynamics and energy barriers of these states (*i*.*e*., sub-states). SMF data provides a quantitative dynamic picture of GPCR activation, which encompasses structural changes on timescales from hundreds of nanoseconds to several milliseconds being controlled by interactions with ligands and intracellular effectors.

## RESULTS

### smFRET reveals ligand-mediated displacement of TM6

smFRET experiments on a double-cysteine mutant (T119C on TM4 and Q226C on TM6) labelled with AF488 (donor) and AF647 (acceptor) show the effect of saturating amounts of four orthosteric ligands on A_2A_R conformations. Observed changes in the FRET efficiency reflect displacements of TM6 relative to TM4, which remains relatively immobile (Supporting Information Section 1, **Figure S1**). Solution smFRET experiments on detergent-reconstituted A_2A_R samples were performed on a custom-built single-molecule fluorescence microscope and the multiparameter photon data was analyzed using custom MATLAB scripts ^35-36^. Labelled protein was diluted to 50 pM in a buffer solution (50 mM HEPES, pH 7.4, 100 mM NaCl, 0.1% MNG-3, 0.02% CHS, 10 mM cysteamine) to ensure a low probability (< 1%) of two independent receptors simultaneously occupying the confocal detection volume. Therefore, nearly all fluorescence intensity bursts originate from single, freely-diffusing, detergent-isolated A_2A_R molecules (**Figure 1a**).

**Figure 1.**
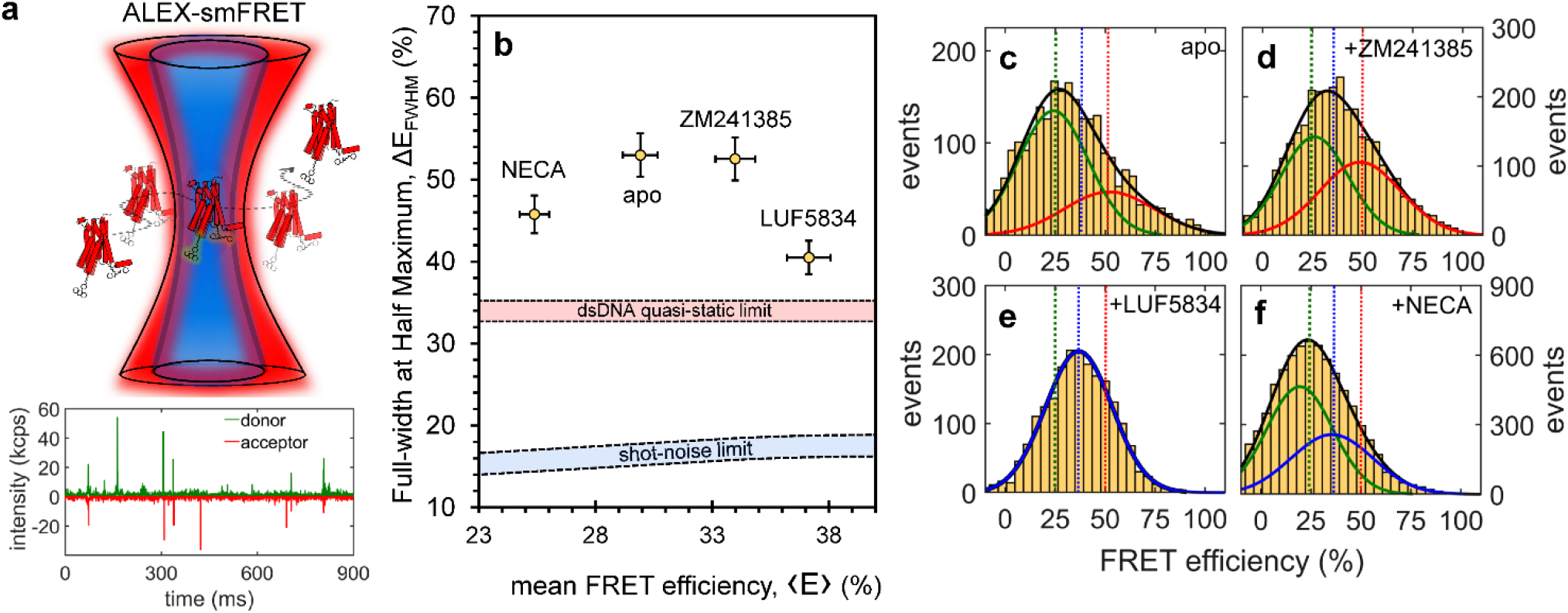
Ligand-induced changes in A_2A_R conformations measured by smFRET. **(a)** A double-cysteine mutant of A_2A_R (T119C Q226C) labelled with AF488 (donor) and AF647 (acceptor) was alternately excited with blue (473 nm) and red (635 nm) lasers; bursts of fluorescence of the donor (**green**) and the acceptor (**red**) were recorded from single receptors diffusing through the confocal detection volume. The FRET efficiency *E* for each burst was calculated using **Eq. 1a** and the values obtained from all bursts for each ligand condition were used to build smFRET histograms **(c-f). (b)** Width (ΔE_FWHM_) *vs*. mean (<E>) for the FRET efficiency distributions shown in **(c-f)**; the error bars were estimated by statistical bootstrapping. Shown as reference are the regions of the shot-noise limit (blue) and of the static limit (dsDNA, pink) (see SI section1.2 for details). **(c-f)** smFRET distributions of A_2A_R in the **(c)** basal state (apo), and in the presence of **(d)** 100 μM ZM241385 (antagonist), **(e)** 100 μM LUF5834 (partial agonist), and **(e)** 100 μM NECA (full agonist), fitted to a sum of Gaussians (black lines). Solid blue, green and red lines represent individual Gaussians and corresponding dashed vertical lines represent their peak center. The results of the fitting are shown in **Table 1**.

smFRET was measured for A_2A_R without ligands (apo) and in the presence of 100 μM ZM241385 (inverse agonist), 100 μM LUF5834 (partial agonist), and 100 μM NECA (full agonist). The corrected energy transfer efficiency *E* for each fluorescence burst was calculated using **Eq. 1**, and all *E* values for each sample were used to build FRET histograms (**Figure 1c-f**). Without any ligands, as well as in the presence of inverse agonist ZM241385, the receptor exhibits relatively broad smFRET distributions centered around ‹*E*› ≈ 30%, with full-width-at-half-maximum (FWHM) values of **ΔE**_**FWHM**_ ≈ 52% (**Figure 1b, Table 1**). The partial agonist LUF5834 and the full agonist NECA shift the smFRET histogram to higher (‹*E*› = 38%) and lower (*E* = 25%) average values, respectively, with concomitant narrowing (**ΔE**_**FWHM**_ = 41% and 46%, respectively, **Table 1**).

**Table 1.** Analysis of smFRET distributions of A_2A_R bound to different orthosteric ligands ^a,b,c^.

| Ligand | $\langle E \rangle$ (%) | $\Delta E_{FWHM}$ (%) | $E_{low}$ (%) | $f_{low}$ | $E_{mid}$ (%) | $f_{mid}$ | $E_{high}$ (%) | $f_{high}$ |
| --- | --- | --- | --- | --- | --- | --- | --- | --- |
| None (apo) | 30 ± 2 | 53 ± 3 | 24 ± 1 | 0.69 ± 0.04 | — | — | 52 ± 1 | 0.31 ± 0.01 |
| +ZM241385 | 34 ± 2 | 52 ± 3 | 26 ± 1 | 0.53 ± 0.02 | — | — | 49 ± 2 | 0.47 ± 0.02 |
| +LUF5834 | 37 ± 2 | 41 ± 2 | — | — | 37 ± 2 | 1.00 | — | — |
| +NECA | 25 ± 1 | 46 ± 2 | 21 ± 3 | 0.51 ± 0.02 | 36 ± 1 | 0.49 ± 0.02 | — | — |

<sup>a</sup> A<sub>2A</sub>R(T119C-G226C) labelled with AF488 and AF647 was measured without (apo) and with different ligands under saturating conditions (i.e., 100 μM).
<sup>c</sup> Mean ( $\langle E \rangle$ ) and width (full-width at half-maximum, $\Delta E_{FWHM}$ ) of smFRET histograms, and the results of fitting (peaks $E_i$ and weights $f_i$ ) to a sum of Gaussians (**Figure 2**)
<sup>c</sup> Standard deviations were estimated by statistical bootstrapping.

To benchmark the conformational dynamics underlying the A_2A_R smFRET histograms, double-stranded DNA (dsDNA) control samples provided a quasi-rigid scaffold for the donor-acceptor pair (Supporting Information Section 1). As shown in **Figure 1b**, the smFRET broadening observed for A_2A_R in all ligand conditions is significantly larger than both the theoretical shot-noise limit computed using the average total number of photons per burst and the quasi-static limit of dsDNA (Supporting Information, section 1.2). While other factors may cause broadening of smFRET histograms (e.g., dye photophysics and orientational freedom), our observations are consistent with low-millisecond exchange between low-FRET and high-FRET states^37^. This range is similar, although somewhat faster than rate of conformational exchange estimated by ^19^F-NMR saturation transfer using different labelling sites ^4, 23^.

Fitting the smFRET histogram for the apo receptor requires two Gaussians, centered at **E**_**low**_ = 24 ± 1% (major) and at **E**_**high**_ = 52 ± 1% (minor) (**Figure 1c**). This corresponds to two states, one where TM6 is relatively farther from TM4 (low FRET, *open/active*) and another where TM6 is relatively closer to TM4 (high FRET, *closed/inactive*). Similar states were resolved previously using smFRET for a different class A GPCR, the β_2_ adrenergic receptor ^26^.

In the presence of the inverse agonist ZM241385, the smFRET histogram is described by the same two FRET states as in the apo form, however the weight of the inactive (high-FRET) state increases (by ∼50%) at the expense of the active (low-FRET) state (**Figure 1d, Table 1**). Binding of the partial agonist LUF5834 leads to a shift of the smFRET histogram to intermediate values between the low-and the high-FRET peaks and to significant narrowing (**Figure 1b**). The histogram is satisfactorily fit with only one Gaussian centered at **E**_**mid**_ = 37 ± 1% (**Figure 1e**). This could originate from accelerated (μs-scale) transitions between the two FRET apo states ^27^, or from a new (partially active) A_2A_R conformation with an intermediate TM6-TM4 separation ^4, 23^ (see also below).

With the full agonist (NECA) bound to receptor, the smFRET histogram exhibits a significant shift towards lower values, but only a small decrease in width compared to the apo state (**Figure 1b**). The FRET distribution is best fitted by the sum of two Gaussians centered at **E**_**low**_ = 21 ± 3 % and **E**_**mid**_ = 36 ± 1 %, having nearly identical weights, *i*.*e*., 0.51 ± 0.02 and 0.49 ± 0.02, respectively (**Figure 1f, Table 1**). These states closely resemble the low-FRET (open/active) state found in the absence of ligands and the intermediate-FRET (partially active) state induced by the partial agonist, respectively. The results of fitting all smFRET histograms, including the error margins, are summarized in **Table 1**.

The Förster radius of the AF488-AF647 dye pair attached to A_2A_R was estimated to be *R*_*o*_ = 50.1 ± 2.5 Å (Supporting Information, section 1.3). As such, using the Förster equation (**Eq. S2**), the donor-acceptor distances (*R*_*DA*_) were estimated to be 49.3 ± 1.4, 54.6 ± 1.6, and 60.6 ± 1.8 Å, for the high, intermediate, and low A_2A_R FRET states, respectively. This corresponds to an ∼10 Å outward movement of TM6 as the receptor transitions from the inactive to active state, which is consistent with crystal structures of A_2A_R in these states (**Figure S1**) ^14-15^. The ligand-dependent shifts and broadening of smFRET distributions suggest that the exchange between active and inactive A_2A_R conformations occurs on timescales that are comparable to the average single-molecule burst duration (2.6 ± 0.5 ms), and much slower than the average inter-photon time in our experiments (∼10 μs). ^37,28^

### BODIPY-FL-labelled A_2A_R: rotational freedom and quenching

To dissect the dynamics of TM6, we labelled A_2A_R at residue V229C with an environment-sensitive dye (BODIPY-FL) with a short linker and performed fluorescence quenching experiments. In the crystal structure of A_2A_R (2YDO/4EIY), several aromatic residues (*i*.*e*., phenylalanine, histidine, and tyrosine) on TM3, TM5 and TM7 are found in the proximity of residue 229 and could act as BODIPY-FL quenchers (**Figure 2a**). Thus, V229C is an adequate construct for monitoring local dynamics in A_2A_R via fluorescence quenching, assuming that quenching dynamics reflects the movement of the host (*i*.*e*., the receptor) and not the intrinsic movement of the dye or its linker ^34^. The linker flexibility and the rotational freedom of BODIPY-FL attached to the apo receptor was measured by fluorescence anisotropy decay (FAD). The anisotropy decay curves were fit to **Eq. 3b** and show that, regardless of the ligand-condition (**Figure S2**), the slow, global rotational dynamics of the receptor (**ρslow** = 52-60 ns) is dominant, while the fast, dye-linker rotational dynamics (**ρfast** = 0.28-0.42 ns) has a minor contribution (∼10 %) to the anisotropy decay (**Figure 2 a-e, Table 2**).

**Table 2.** Time-resolved fluorescence anisotropy parameters of A_2A_R(V229C)-BODIPY-FL in different ligand states ^a,b,c^

|  | Apo | +ZM241385 | +LUF5834 | +NECA |
| --- | --- | --- | --- | --- |
| $\rho_{fast}$ (ns) | 0.42 ± 0.04 | 0.28 ± 0.04 | 0.36 ± 0.03 | 0.42 ± 0.04 |
| $\rho_{slow}$ (ns) | 58 ± 6 | 60 ± 4 | 52 ± 5 | 60 ± 6 |
| $A_{fast}$ | 0.09 ± 0.01 | 0.10 ± 0.01 | 0.09 ± 0.01 | 0.09 ± 0.01 |
| $A_{slow}$ | 0.91 ± 0.01 | 0.90 ± 0.01 | 0.91 ± 0.01 | 0.91 ± 0.01 |

<sup>a</sup> Measurements were performed on solutions of 100 nM BODIPY-FL-labelled receptors without (apo) and with saturating amounts (100 μM) of ligands
<sup>b</sup> The experimental data were fitted to multi-exponential decay model (**Eq. 3b**)
<sup>c</sup> Standard deviations were estimated by parametric bootstrapping.

**Figure 2.**
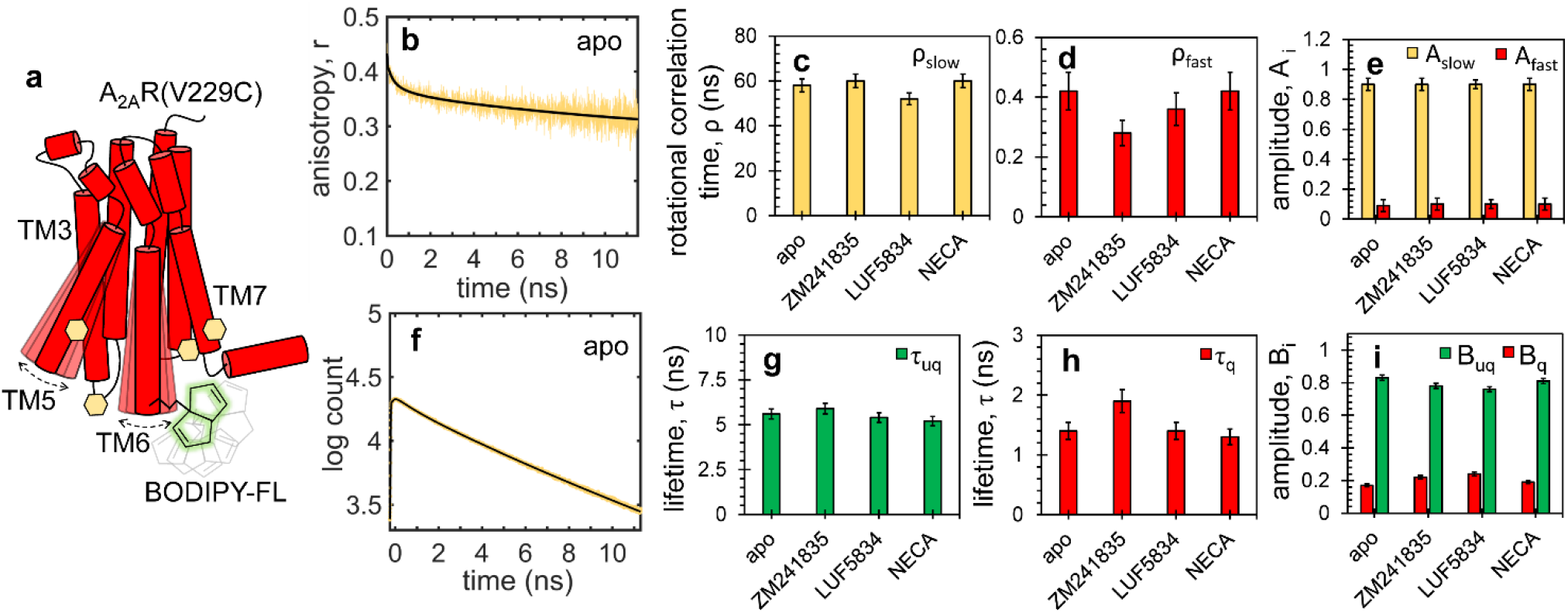
Rotational freedom and quenching of BODIPY-FL in A_2A_R. **(a)** Native aromatic residues (yellow hexagons) on TM3, TM5, and TM7 provide a dynamic quenching environment for BODIPY-FL (green) attached to residue 229 in A_2A_R(V229C). Fluorescence anisotropy **(b)** and lifetime **(f)** decay curves for apo A_2A_R (yellow) were fitted to multi-exponential functions, i.e. **Eq. 3b** and **Eq**.**4**, respectively (black). Similar data was acquired under excess (100 μM) of ligands, i.e. inverse agonist ZM241385, partial agonist LUF5834 and full agonist NECA, and was fitted to the same models. Slow and fast rotational correlation times for all ligand conditions shown as yellow and red bars in bars in **(c)** and **(d)**, respectively, and their corresponding amplitudes **(e)**. Unquenched and quenched fluorescence lifetimes shown as green and red bars in **(g)** and how **(h)**, respectively, and their corresponding amplitudes **(i)**. Standard deviations shown in **(c-e)** and **(g-i)** were estimated by parametric bootstrapping. The experiments were performed on 100 nM solutions of labelled receptors using pulsed 480-nm laser excitation. The fitting results are summarized in **Table 2** (anisotropy) and **Table 3** (lifetime).

Fluorescence lifetime measurements describe the extent of quenching when BODIPY-FL is attached to residue 229 in A_2A_R (**Figure 2 f-i, Table 3)**. For the apo receptor (**Figure 2f**), the experimental decay curve was best fitted to a sum of two exponential lifetimes, **τ**_**q**_ = 1.4 ± 0.1 ns and **τ**_**nq**_ = 5.6 ± 0.1 ns, with amplitudes **B**_**q**_ = 0.17 ± 0.01 and **B**_**nq**_ = 0.83 ± 0.01, respectively (**Eq. 4**). The long lifetime is similar to previous values reported for free BODIPY-FL in solution (*i*.*e*., 5.5 ns) ^38^, while the short lifetime corresponds to the quenched state induced by the host protein. The two lifetimes and amplitudes vary slightly across different ligand conditions, with most quenching occurring in the presence of partial and inverse agonists. Thus, the fluorescence of BODIPY-FL attached to TM6 at residue 229 is partially quenched, and the quenching varies in response to different ligand-induced rearrangements of TM bundles or local helical movements.

**Table 3.** Fluorescence lifetime parameters of A_2A_R(V229C)-BODIPY-FL in different ligand states ^a,b,c^

|  | Apo | +ZM241385 | +LUF5834 | +NECA |
| --- | --- | --- | --- | --- |
| $\tau_{nq}$ (ns) | 5.6 ± 0.1 | 5.9 ± 0.2 | 5.4 ± 0.1 | 5.2 ± 0.2 |
| $\tau_q$ (ns) | 1.4 ± 0.2 | 1.9 ± 0.2 | 1.4 ± 0.2 | 1.3 ± 0.2 |
| $B_{nq}$ | 0.83 ± 0.1 | 0.78 ± 0.01 | 0.76 ± 0.01 | 0.81 ± 0.01 |
| $B_q$ | 0.17 ± 0.01 | 0.22 ± 0.01 | 0.24 ± 0.01 | 0.19 ± 0.01 |

<sup>a</sup> Measurements were performed on solutions of 100 nM BODIPY-FL-labelled receptors without (apo) and with saturating amounts (100 μM) of ligands
<sup>b</sup> The experimental data were fitted to multi-exponential decay model (**Eq. 4**)
<sup>c</sup> Standard deviations were estimated by parametric bootstrapping.

### BODIPY-FL as a probe for PET-FCS

To benchmark the intrinsic photoblinking of BODIPY-FL due to reversible transitions to its dark (triplet) state, FCS measurements of the free dye in solution were performed under the same buffer conditions as the receptor (**Figure S3a, Table S1**), including photo-protectants (20 mM cysteamine and 2 mM Trolox) to enhance brightness and reduce photoblinking ^39^. The measured autocorrelation curve is best fitted to a diffusion model with one triplet component (**Eq. 5**, with no quenching terms), and the triplet lifetime obtained (**τ**_**t**_ = 3 ± 1 μs) is in agreement with previous reports (2-8 μs) ^40^. For the global analysis of FCS data of the receptor under different ligand conditions, the BODIPY-FL triplet lifetime was used as a shared fitting parameter (**Eq. 5**, see also below).

Of the several aromatic residues in A_2A_R near residue 229, tyrosines are most likely to act as PET quenchers of BODIPY-FL fluorescence, due to their abundance and quenching power ^30^. To verify the quenching of BODIPY-FL by tyrosine, FCS experiments of free dye (∼10 nM) were performed in the presence of 1 mM and 3 mM tyrosine under similar buffer conditions as above (**Figure S3b-c**). Data was fitted to **Eq. 5** and the fitting parameters are summarized in **Table S1**. Fitting of the FCS curve obtained upon adding 1 mM tyrosine requires an additional lifetime (∼40 μs) compared to the free dye in buffer solution. Upon increasing the tyrosine concentration to 3 mM, this decay component becomes more pronounced and also a faster decay component (∼100 ns) appears.

Quenching collisions corresponding to the intermediate PET lifetime (∼40 μs) occur with a frequency of ∼25 ms^-1^, whereas those responsible for the fast PET lifetime (∼100 ns) occur with a frequency of ∼10^5^ ms^-1^. These rates represent diffusion-limited quenching interactions between BODIPY-FL and tyrosine in an aqueous solution ^41^. In A_2A_R, the effective concentration of tyrosine in the proximity of the dye is higher (∼60 mM), which would normally lead to an increase of the quenching rates. However, the PET quenching rates in A_2A_R will be determined by both local helical dynamics and long-range TM movements, which can alter the proximity and orientation of BODIPY-FL relative to native A_2A_R tyrosine residues ^34^.

### Direct measurements of A_2A_R conformational dynamics by PET-FCS

PET-FCS measurements were performed on the apo receptor, and upon addition of 100 μM of inverse agonist, partial agonist, or full agonist, as well as in the presence of 10 μM mini-G_s_ and 100 μM NECA, 2 mM Mg^2+^ and 50 μM GDP (**Figure 3**). At the concentration of BODIPY labelled-A_2A_R used in FCS experiments (∼10 nM) there is an approximately 10^4^ -fold excess of ligand and 10^3^-fold excess of mini-G_s_. For clarity, the fitted diffusion component was subtracted from each experimental FCS curve to obtain decay curves that included only contributions from photoblinking of the dye (*i*.*e*., triplet and PET quenching), as illustrated in **Figure 3a** and described previously ^34^. The autocorrelation curves obtained for each sample condition were fitted to **Eq. 5** and the results are summarized in **Table 4**. Using the diffusion times and the Stokes-Einstein equation (**Eq. 6)**, the hydrodynamic radii were estimated to be **R**_**H**_ = 6 ± 1 nm for all samples (slightly higher for apo, 7 ± 1 nm), which matches the size of the detergent micelle encapsulating the receptor. The data was fitted to a model including one triplet decay and three PET quenching components. Global fitting of all FCS curves was performed, with the triplet lifetime shared and close to that of the free dye (**t**_**trip**_ = 5 ± 1 μs, **Table 4**), and with ligand-specific PET lifetimes and amplitudes.

**Table 4.**
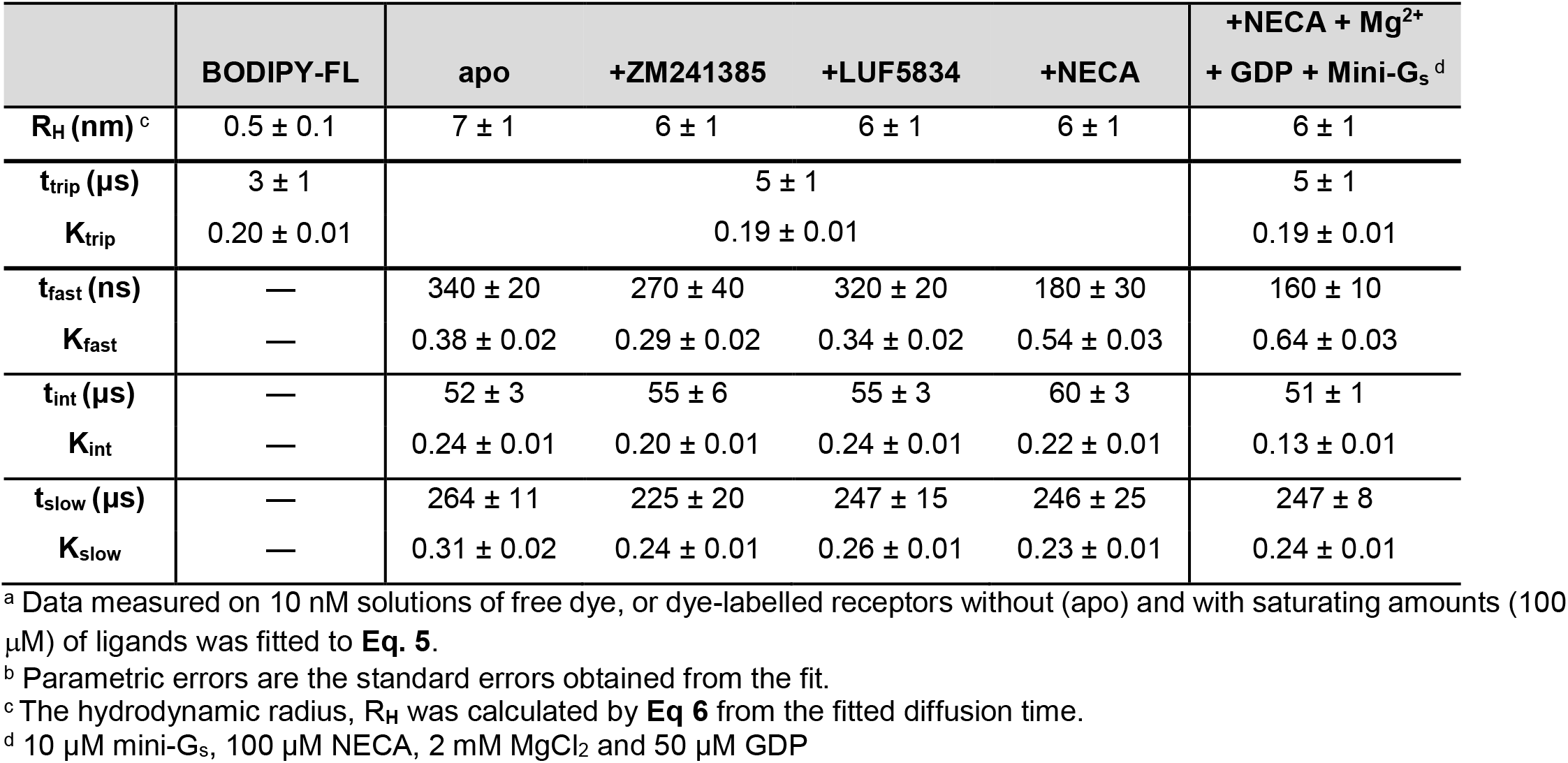
FCS fitting parameters for free BODIPY-FL and for BODIPY-FL-labelled A_2A_R bound to different ligands ^a,b^

|  | BODIPY-FL | apo | +ZM241385 | +LUF5834 | +NECA | +NECA + Mg<sup>2+</sup><br>+ GDP + Mini-G<sub>s</sub> <sup>d</sup> |
| --- | --- | --- | --- | --- | --- | --- |
| **R<sub>H</sub> (nm)** <sup>c</sup> | 0.5 ± 0.1 | 7 ± 1 | 6 ± 1 | 6 ± 1 | 6 ± 1 | 6 ± 1 |
| **t<sub>trip</sub> (μs)** | 3 ± 1 | 5 ± 1 |  |  |  | 5 ± 1 |
| **K<sub>trip</sub>** | 0.20 ± 0.01 | 0.19 ± 0.01 |  |  |  | 0.19 ± 0.01 |
| **t<sub>fast</sub> (ns)** | — | 340 ± 20 | 270 ± 40 | 320 ± 20 | 180 ± 30 | 160 ± 10 |
| **K<sub>fast</sub>** | — | 0.38 ± 0.02 | 0.29 ± 0.02 | 0.34 ± 0.02 | 0.54 ± 0.03 | 0.64 ± 0.03 |
| **t<sub>int</sub> (μs)** | — | 52 ± 3 | 55 ± 6 | 55 ± 3 | 60 ± 3 | 51 ± 1 |
| **K<sub>int</sub>** | — | 0.24 ± 0.01 | 0.20 ± 0.01 | 0.24 ± 0.01 | 0.22 ± 0.01 | 0.13 ± 0.01 |
| **t<sub>slow</sub> (μs)** | — | 264 ± 11 | 225 ± 20 | 247 ± 15 | 246 ± 25 | 247 ± 8 |
| **K<sub>slow</sub>** | — | 0.31 ± 0.02 | 0.24 ± 0.01 | 0.26 ± 0.01 | 0.23 ± 0.01 | 0.24 ± 0.01 |

<sup>a</sup> Data measured on 10 nM solutions of free dye, or dye-labelled receptors without (apo) and with saturating amounts (100 μM) of ligands was fitted to **Eq. 5**.
<sup>b</sup> Parametric errors are the standard errors obtained from the fit.
<sup>c</sup> The hydrodynamic radius, R<sub>H</sub> was calculated by **Eq 6** from the fitted diffusion time.
<sup>d</sup> 10 μM mini-G<sub>s</sub>, 100 μM NECA, 2 mM MgCl<sub>2</sub> and 50 μM GDP

**Figure 3.**
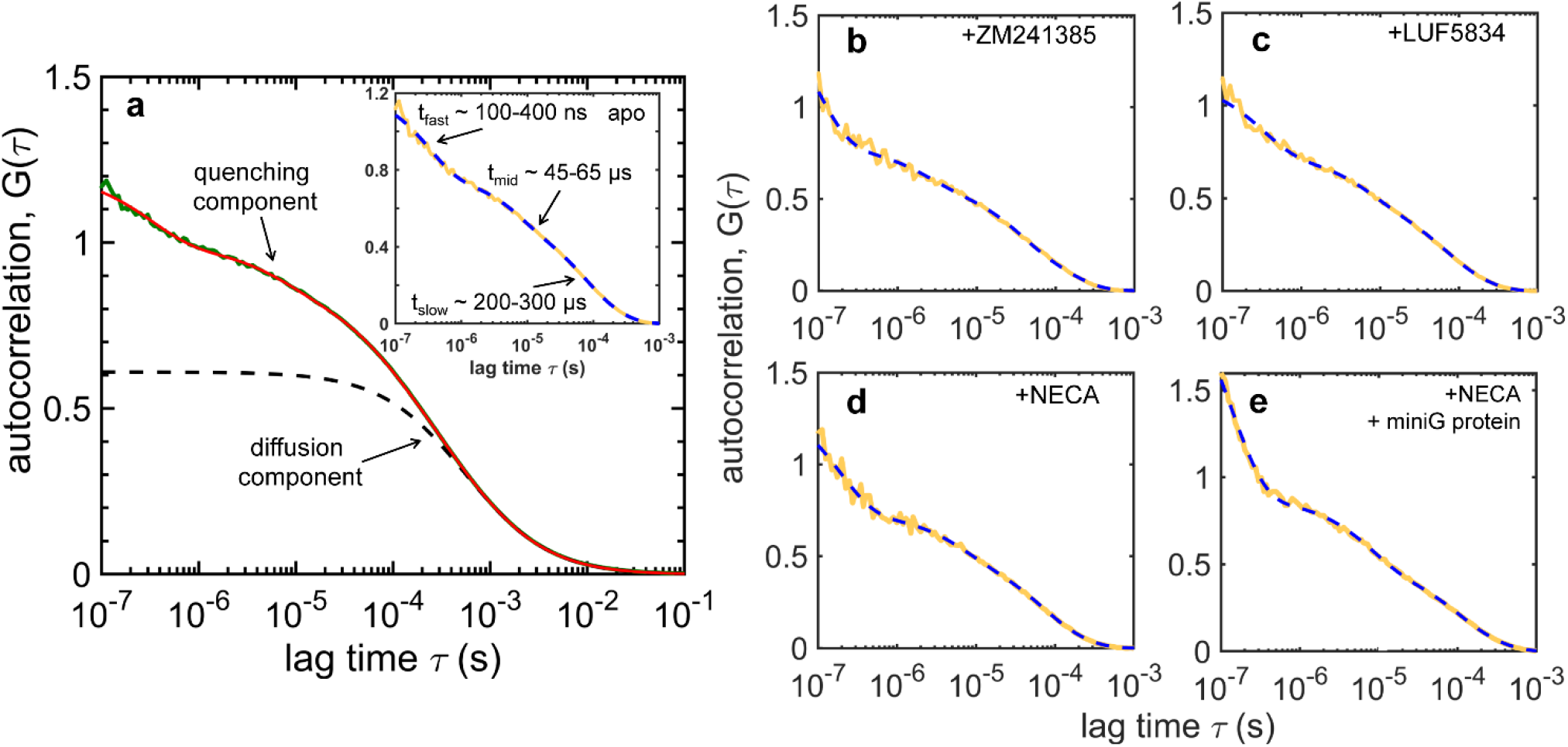
FCS measurements of conformational dynamics around residue 229 in A_2A_R. **(a)** FCS data (green line) of A_2A_R-V229C-BODIPY-FL fitted to **Eq**.**5** (red line). The pure diffusion component in the fitted curve (black dashed line) can be subtracted from the data to highlight the decays due to triplet and quenching. The inset shows the curves obtained after removing the diffusion term from the raw data (solid yellow) and the fitted curve (dashed blue). “Diffusion-free” FCS curves of the labelled receptor in the presence of **(b)** 100 μM ZM241385, **(c)** 100 μM LUF5834, **(d)** 100 μM NECA, and **(e)** 100 μM NECA, 2 mM MgCl_2_, 50 μM GDP and 10 μM mini G protein, respectively; raw data (solid yellow) and fitted curves to **Eq. 5** (dashed blue). The FCS fitting results for all sample conditions are summarized in **Table 4**.

The ligand-specific PET quenching lifetimes can be grouped into *fast* (**t**_**q, fast**_ = 150-350 ns), *intermediate* (**t**_**q, int**_ = 50-60 μs), and *slow* (**t**_**q, slow**_ = 200-300 μs), with corresponding amplitudes (***K***_***i***_) that vary as a function of ligand (**Figure 4a-f, Table 4**). Most notably, the contribution of the fast kinetics increases in the fully active states (NECA-and mini-G_s_-bound) relative to those from two slower kinetic components, as shown by the amplitudes ***K***_***i***_ (**Figure 4d-f)**. The fast kinetics becomes even faster in the active states, with its lifetime decreasing from 340 ± 20 ns in the apo state to 180 ± 30 ns in the NECA-bound state and to 160 ± 10 ns in the mini-G_s_-bound state (**Figure 4a**). The intermediate lifetime slightly increases upon activation by NECA, *i*.*e*., from 52 ± 3 μs to 60 ± 3 μs, but it returns to the apo-state level (51 ± 1 μs) upon binding of mini-G_s_ (**Figure 4b**). Similarly, the lifetime of the slow component is nearly unchanged between the apo and the partially/fully active states (245-265 μs), and it shows only a shows a minor decrease in the inverse-agonist-bound state (225 ± 20 μs) (**Figure 4c**).

**Figure 4.**
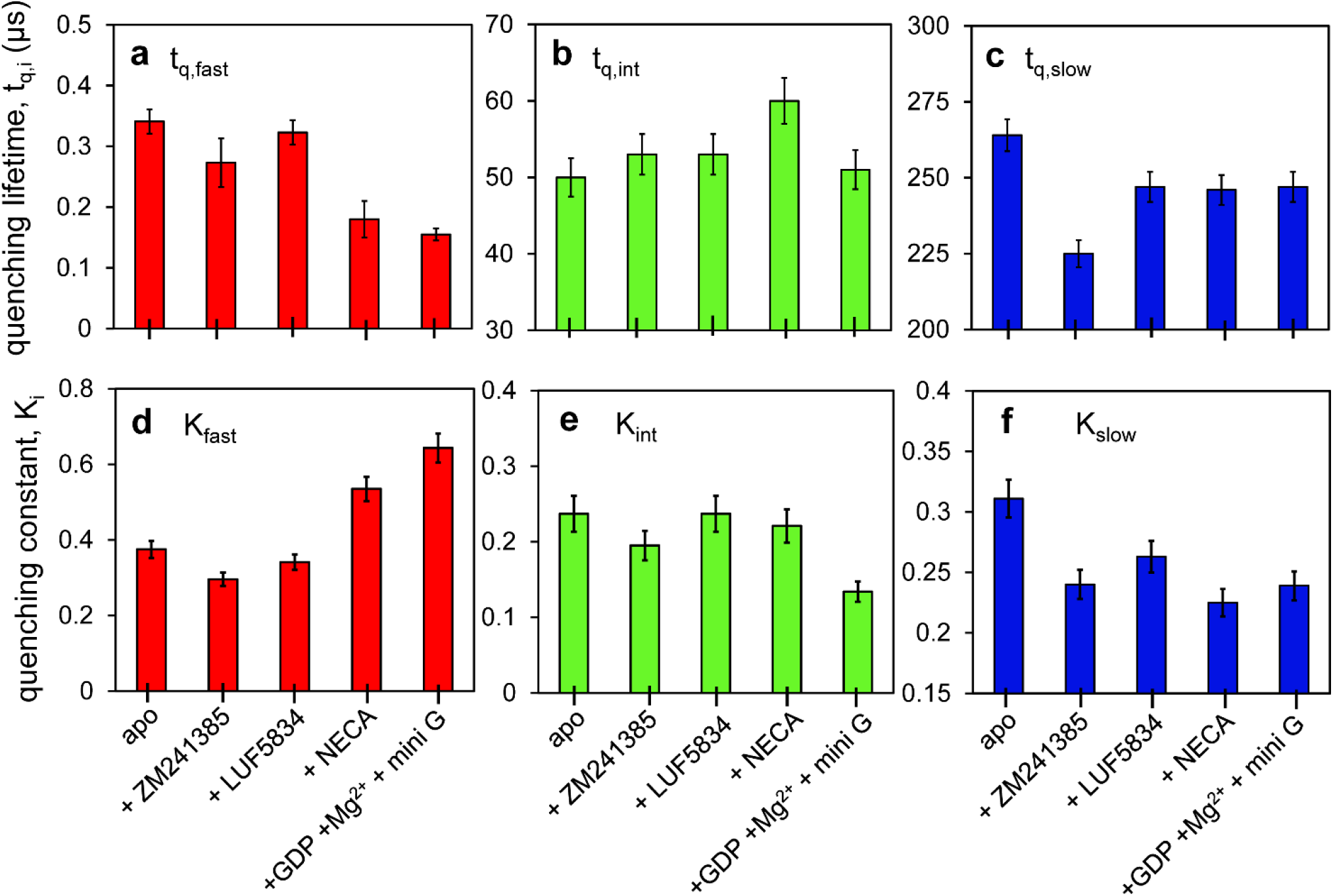
PET quenching lifetimes and amplitudes in A_2A_R as modulated by ligands and a G-protein mimetic. PET lifetimes (**t**_**q**,**i**_) and corresponding amplitudes **(K_i_)** obtained by fitting FCS data shown in **Figure 3** to a model described by **Eq. 5**. Three PET components were resolved for each ligand condition: *fast* **(b, d)** *intermediate* (**c, e)**, and *slow* **(d, f)**, with varying lifetimes and amplitudes. The error bars were derived by parametric bootstrapping. The fitting results are summarized in **Table 4**.

The intermediate and the slow PET lifetimes (30-300 μs) reflect changes in the brightness of the dye that are likely driven by local helical movements and rearrangement of the TM bundles, as suggested previously ^34^. These timescales of motion are on the same order of magnitude as those suggested by exchange broadening of the smFRET distributions (**Figure 1**, see above). In particular, the microsecond protein dynamics captured by PET-FCS could be associated to the on/off behavior of the ionic-lock for the salt-bridge between R102 of TM3 and E228 of TM6 ^4, 40^.

### Molecular dynamics simulations of PET quenching in A_2A_R

MD simulations of A_2A_R were performed to identity the native aromatic residues and the protein motions responsible for the PET quenching dynamics revealed by FCS. To compute the spatial extension of the BODIPY-FL dye from its anchoring point on the receptor (**Figure S4a**,**b**), we simulated an 11-residue segment of TM6 centered at residue 229, with the dye attached at this residue (**Figure 5a**). The dye linker relaxes rapidly, with an exponential distance autocorrelation lifetime of *τ*_acor_ = 0.8 ± 0.3 ns (**Figure S4c**).

**Figure 5.**
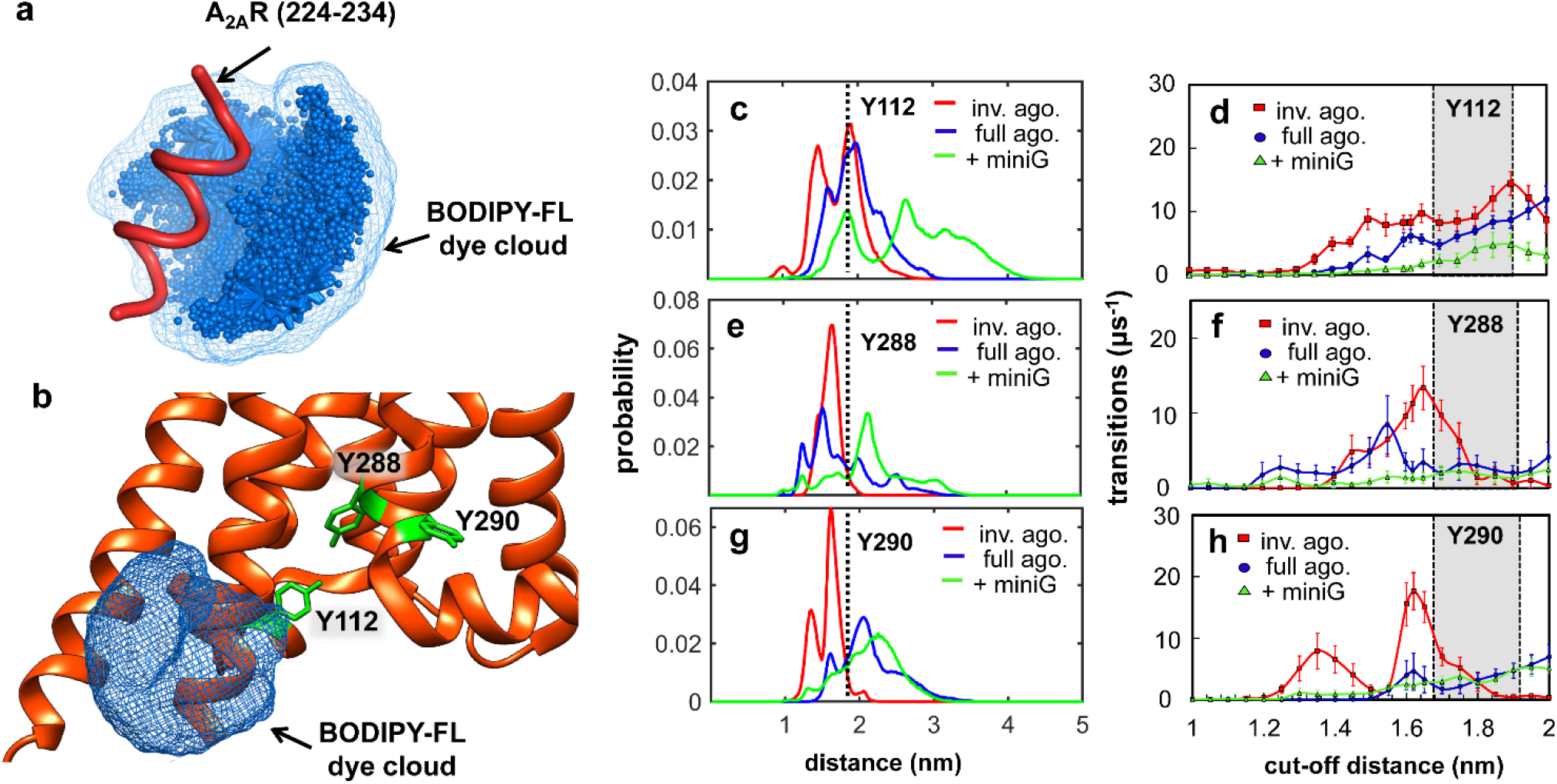
Molecular dynamics simulations of PET quenching dynamics in A_2A_R. **(a)** Simulations of the cloud of possible positions of a BODIPY-FL molecule (with boron replaced by carbon) attached to position 229 within a small peptide segment of A_2A_R (224-234). **(b)** The simulated BODIPY-FL cloud from **(a)** overlaid with the PDB structure of A_2A_R, with tyrosine residues (green) in the proximity representing possible PET quenching candidates. Probability distributions for the distance between the alpha-carbon of position 229 to the center of mass of the aromatic system of tyrosine Y112 on TM3 **(c)**, Y288 **(e)** and Y290 **(g)** on TM7: inverse agonist (red), full agonist (blue), full agonist + miniG (green). Inter-residue distance corresponding to the cut-off distance for PET quenching of BODIPY-FL by tyrosine, *d*_*q*_ = 1.81 nm, is shown as a vertical dash line. The frequency of quenching/de-quenching transitions for Y112 **(d)**, Y288 **(f)** and Y290 **(h)** as a function of cut-off distance in presence of inverse agonist (red squares), full agonist (blue circles), and full agonist + mini G (green triangles).Error bars are standard deviations from 6 replicate simulations. The shaded regions in **(d**,**f**,**h)** represent the most likely cut-off distance for PET quenching of BODIPY-FL by tyrosine, *d*_*q*_ = 1.8 ± 0.1 nm. MD simulations were initiated using crystal structures in which the A_2A_R was crystalized bound to an inverse agonist, an agonist, or an agonist plus mini-G, as shown in **Table S2**.

This rapid relaxation permitted a simplifying approximation in which simulations of unmodified A_2A_R were post-processed to mimic the presence of a temporally averaged dye cloud (**Figure 5a**,**b**). In order to quantify the dynamic quenching of BODIPY-FL by specific aromatic residues in the receptor, we defined the radius of a spherical surface surrounding the C_β_ atom of A_2A_R residue V229 as <*l>* = 0.81 ± 0.01 nm. This value represents the mean distance from the C_β_ atom of V229 to the center of geometry of non-hydrogen atoms in the dye’s aromatic group obtained from three replicate simulations (**Figure S4**). Since the typical quenching distance for BODIPY-FL is 1.0 nm or less ^29, 42^, we defined a quenching cutoff distance of *d*_q_ = (1+ <*l>*) = 1.81 nm from the C_β_ atom of V229 to the center of mass of the side chain of each quenching candidate residue in A_2A_R (**Figure S5**).

Simulations of wild-type apo A_2A_R in a hydrated lipid bilayer composed of 1-palmitoyl-2-oleoyl-*sn*-glycero-3-phosphocholine (POPC) were performed. An aggregate time of 210 μs was obtained via a total of 42 independent 5-μs simulations initiated from seven different crystal structures (see **Table S2**). Since our simulations were conducted with the wild-type unconjugated receptor, we quantified quenching based on the distance, *d*, from the C_β_ atom of V229 to the center of mass of each aromatic side chain in the simulated region of the A_2A_R (S6 to V307). Given the “dye-cloud” approximation (**Figure 5a**,**b**), we applied a cutoff distance of *d*_q_ = 1.8 ± 0.1 nm, such that states with *d* < *d*_q_ are defined to be “quenched” (dark) and states with *d* > *d*_q_ are defined to be “unquenched” (fluorescent) (**Figure S5**). The inter-residue distance distributions from the C_β_ atom of V229 to all possible quenching candidates (Trp, Tyr, Phe and His) show that 6 of the 42 residues monitored have overall populations that span across the *d*_q_ cutoff (dashed red arrows, **Figure S6**): Y103, Y112, Y197, Y288, Y290, and F295 (see e.g., **Figure 5 c,e,g**).

To quantify reversible transitions between quenched and unquenched states, we analyzed the time trajectories of *d* in each simulation, separately for each of the 6 residues above. Statistical noise was reduced by smoothening the trajectories with a 20-ns running average and transitions between quenched and unquenched states were interpreted using a cut-off distance that was varied between *d*_q_ = 1 nm and *d*_q_ = 2 nm. By comparing the frequency of MD quenching/dequenching transitions to the experimental values, we propose that Y112 on TM3, and Y288 and Y290 on TM7 are likely responsible for the fast PET-FCS lifetime. These tyrosines exhibit a maximum number of quenching-dequenching transitions at the expected cut-off distance of 1.8 ± 0.1 nm for each ligand condition (**Figure 5 d**,**f**,**h**). The MD estimates of the PET rates are an order of magnitude higher than the experimental values and should be viewed as an upper limit, as the relative orientation of the dye and the tyrosine, which is critical for electron transfer, has not been taken into account in the calculations.

The residue Y112 is on an alpha-helical domain on TM3 and forms a helical bridge to TM4, which makes frequent transitions (*i*.*e*., 4–10 μs^-1^) from an ordered helical configuration to a disordered loop-like configuration in the MD simulations (**Figure 5d**). This transition to a loop-like configuration brings Y112 within quenching range of BODIPY-FL on residue V229 (**Movie S1**) and is more pronounced in the inactive state (red lines, **Figure 5c**,**d**). Comparing the MD-estimated quenching/dequenching rate to that estimated by PET-FCS in the inverse-agonist-bound state (*i*.*e*., 1/ ***t***_**q**,**fast ∼**_ 3.7 μs^-1^), the fast PET lifetime can be assigned to quenching by Y112. As the receptor switches into more active states, such as those stabilized by full agonist and mini-G protein, however, the frequency of Y112 transitions decreases, implying stabilization of the helix (blue, and green lines, **Figure 5c**,**d**). This decreasing trend in PET dynamics can be rationalized in terms of the crystal structure of the mini-G-bound receptor (5G53), which shows stabilizing interactions between Y112 of TM3 of A_2A_R and H387 of the mini-G protein ^43^ (see Supporting Information, **Figure S7a**,**b**).

The other two candidates, Y288 and Y290 on TM7 show rather opposing trends in MD simulations (**Figure 5e**,**g**,**f**,**h**), such that the inverse-agonist bound state exhibits less frequent quenching transitions (∼1.6 μs^-1^) when compared to the full-agonist-bound (∼3.0 μs^-1^), or to the receptor bound to full-agonist and mini-G-protein (∼1.9 μs^-1^) (**Figure 5f**,**g**). The increasing number of transitions estimated by MD for Y288 and Y290 upon receptor activation suggests that there are larger fluctuations in the active states of A_2A_R that bring these two aromatic residues on TM7 within quenching range of BODIPY-FL attached to TM6. Another candidate quencher residue, Y197 on TM5, shows little to no transitions in the presence of inverse and full agonist, and it is only significant in simulations initiated with a receptor that is pre-organized to bind the mini-G protein. This suggests a dynamic reversible rotation of TM5 relative to TM6 in a manner that increases the accessibility of Y197 to residue V229 (**Figure S7c-e**).

## DISCUSSION

This study combines fluorescence spectroscopy and molecular dynamics to examine the conformational dynamics of A_2A_R over a wide range of timescales. A_2A_R conformations were measured at the single-molecule level using FRET between specific sites in TM4 and TM6 that were labelled with a donor-acceptor dye pair. The smFRET data confirm previous ^19^F-NMR findings that active (*i*.*e*., S_3_ and S_3’_) and inactive (*i*.*e*., S_1-2_) states of the receptor co-exist and ligands alter their relative population ^4-5^. Specifically, the broadening of the smFRET histogram for the apo receptor is consistent with the existence of different FRET sub-populations and millisecond exchange between high-FRET inactive (S_1_ and S_2_) and low-FRET active (S_3_ and S_3’_) A_2A_R conformations. In addition, using the quasi-static dsDNA reference, each FRET sub-population appears to be broadened (**ΔE**_**FWHM-low**_ ∼ 40% and **ΔE**_**FWHM-high**_ ∼ 50%, **Figure 1**), suggesting millisecond-scale exchange between S_3_ and S_3’,_ and between S_1_ and S_2_, respectively. Previous ^19^F-NMR studies have conflicting outcomes regarding the basal activity of A_2A_R, reporting either a high ^4^ or a low ^23^ fraction of active or activation intermediate states for the apo receptor. Our smFRET data confirms that the ligand-free A_2A_R is inherently dynamic, with a population bias towards active-like conformations that is consistent with significant basal activity ^44^.

Binding of the inverse agonist ZM241385 did not change the overall broadening of the smFRET distribution, but led to ∼50% increase in the fraction of inactive (high-FRET) sub-population (**Table 1**). This is consistent with a ^19^F-NMR study that showed a similar reequilibration of the apo conformational ensemble in the presence of inverse agonist ^23^. An alternative interpretation would invoke the emergence of an intermediate state between the active and inactive states (dashed blue line between the green and red lines, **Figure 1d**), corresponding to the S_3_ state that is enriched in the inverse-agonist-bound receptor ^4^.

The partial agonist LUF5834 shifts the smFRET distribution to intermediate values between the low-and high-FRET sub-populations (**Figure 1e**). It also considerably narrows the distribution to a level that is close to the quasi-static control (dsDNA) for smFRET. This is consistent with the partial agonist stabilizing a distinct A_2A_R conformation (blue line, **Figure 1e**), which corresponds to the S_3_ state identified by ^19^FNMR ^4^. Alternatively, the narrow smFRET peak could arise from a significantly increased exchange rate between the apo low-and the high-FRET states (∼1-10 μs lifetime), although this appears to be at least 2 orders of magnitude faster than the exchange rates inferred by 2D [^19^F,^19^F]-EXSY ^23^.

The full agonist NECA shifts the smFRET distribution to lower *E* while maintaining significant broadening (**Figure 1f**). The data analysis reveals two nearly equal sub-populations that are similar to the low-FRET conformation of the apo receptor, and the mid-FRET conformation stabilized by partial agonist. smFRET confirms previous ^19^FNMR observations of two A_2A_R active states for the NECA-bound receptor ^4, 23^, with blue and green lines in **Figure 1f** corresponding to S_3_ and S_3’_, respectively. Overall, the smFRET results provide new, independent evidence that inactive and active A_2A_R conformations co-exist in the absence of ligands, that transitions between these states occur on a millisecond timescale, and that ligand binding alters both the energetics and the dynamics of these states.

In previous ^19^FNMR studies, the point mutation V229C on TM6 provided good sensitivity to discriminate between active and inactive states, while still maintaining the overall integrity and function of A_2A_R ^4-5^. These studies confirmed the co-existence of inactive and active states in the apo receptor; however, direct measurements of (fast/microsecond) conformational dynamics is still missing. To fill this gap, we performed fluorescence quenching experiments on A_2A_R-V229C labelled with BODIPY-FL in the absence and presence of ligands (**Figures 2-4**). PET is a dynamic (collisional) quenching process, in which the formation a weak complex between an organic dye and a quencher molecule results in the transfer of an excited electron from the donor (dye) to the acceptor (quencher) molecule causing a reduction in the excited-state lifetime of the dye ^31^. Electron transfer is a short-range process occurring at dye-quencher distances less than ∼10 Å and for favorable orientations between the two molecules. As A_2A_R has several aromatic residues in the vicinity of residue 229, the displacement and orientation of TM6 will determine the proliferation of PET quenching interactions of the probe.

A short two-carbon linker between BODIPY-FL and the point of attachment on A_2A_R (V229C) ensures that quenching of the dye is primarily due to the movement of TM6 (relative to tyrosines on TM3, TM5, and TM7) and not due to the intrinsic movement of the dye/linker. Indeed, fluorescence lifetime and anisotropy decay analysis (**Figure 2, Table 2**) reveal that quenching occurs upon labelling the receptor, and that the dye reorients slowly, on a timescale determined by the global rotation of the receptor. Additionally, the smFRET data (see above) shows binding of partial/full agonist displaces TM6 by ∼ 5-10 Å relative to TM4, which is consistent with the reduction in distance required for quenching to occur between BODIPY-FL and tyrosine residues on TM7 (Y288 and Y290).

We made inferences on the (sub)microsecond dynamics of A_2A_R based on PET quenching lifetimes (**t**_**q**_) measured by FCS. As such, fluorescence intensity fluctuations of BODIPY-FL-labelled A_2A_R were found to occur on *fast* (**t**_**q, fast**_ = 150-350 ns), *intermediate* (**t**_**q, int**_ = 50-60 μs) and *slow* (**t**_**q, slow**_ = 200-300 μs) timescales, and they were modulated by orthosteric ligands (**Figure 4, Table 4**). These lifetimes represent timescales of the local re-arrangement of TM3, TM5, TM6, and TM7, which either proliferate or suppress quenching interactions between BODIPY-FL and tyrosine residues located on these TMs^34^.

Using MD simulations, we identified the residue Y112 on intracellular loop 2 (ICL2) between TM3 and TM4 as a promising candidate for dynamic quenching of BODIPY-FL on TM6 (**Movie S1**). Nevertheless, whereas PET-FCS indicates that addition of full agonist and mini-G_α_ significantly increases the frequency of reversible quenching (**Figure 4a**,**d**), MD simulations predict fewer quenching transitions of Y112 as A_2A_R trends towards full activity (**Figure 5c**,**d**). One possible explanation for this discrepancy could be the polarization bias in FCS measurements.^34^ However, the FAD-measured rotational correlation time of A_2A_R is 50-60 ns (**Figure 2**), i.e., 3-5 times faster than **t**_**q, fast**_ in FCS. Thus, the fastest FCS decay predominantly reports on the dynamics of PET quenching, not depolarization. Corroborating this, FCS experiments on the free dye in a solution of 3 mM tyrosine reveal a similar quenching lifetime (**t**_**q**_ ∼100 ns, **Figure S3**) as in A_2A_R, while the depolarization of the dye in solution is much faster (**ρ** ∼ 0.3 ns, **Table 2**).

Therefore, we propose that conformational changes involved in receptor activation bring other aromatic quenchers into the proximity of BODIPY-FL. In MD simulations, residues Y288 and Y290 exhibit distance fluctuations to and from residue 229 which significantly increase as the receptor trends toward full activity (**Figure 5 e-h**). These fluctuations bringing Y288 and Y290 on TM7 within close proximity to residue 229 on TM6 may be responsible for the increase in the rate and the amplitude of fast PET-FCS component for the full agonist and mini-G protein bound case (**Figure 4a**,**d**). Another residue on TM5, Y197, may have a negligible contribution to the fast quenching phase in the absence of ligands, however it contributes significantly in the presence of mini-G (**Figure S7d**,**e**).

The sub-millisecond dynamics reveled by PET-FCS could also be related to the integrity of the ionic lock between R102 (TM3) and E228 (TM6), which was shown to be altered by different ligands ^4-5^. 2D probability plots were generated to assess the integrity of the ionic lock in the inactive (inverse agonist), active (full agonist), and full agonist plus mini-G protein bound structures, respectively (**Figure S8**). As suggested previously, when the minimum distance between the two residues, 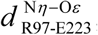, is less than 5 Å, the ionic lock remains intact due to formation of a salt bridge, whereas at longer distances the ionic lock is broken ^45-46^. In simulations with the inverse agonist, the ionic lock is mostly intact (**Figure S8a**), however, in simulations initiated from agonist (**Figure S8b**) or mini-G (**Figure S8c**) bound configurations, the ionic lock is predominantly open.

For the active structures, two populations emerge, conformations where TM3 and TM6 are close to each other (∼3-5 Å, ionic lock intact) or far apart (∼5 Å, ionic lock broken), with a significant population in the latter (**Figure S8b**,**c**). The breaking of this ionic lock may proliferate the movement of TM3 or TM6 in a manner that causes (sub) microsecond fluctuations in local helical regions or in the TM domains at large.

## CONCLUSIONS

Single-molecule fluorescence techniques were applied to the A_2A_ Adenosine Receptor and revealed new details about its structural plasticity and how this is modulated by ligands and a G-protein mimetic. smFRET analysis showed that orthosteric ligands alter the relative population of inactive and active conformations, with apo A_2A_R biased towards active states, and the inverse agonist increasing the fraction of inactive states. The partial agonist stabilizes an intermediate conformation with a TM6-TM4 separation distance that lies between the active (open/high-FRET) and inactive (closed/low-FRET) states. The full agonist promotes an active conformation of the receptor, which coexists with a nearly equally populated intermediate conformation. The broadening of smFRET distributions implies low-millisecond (>3ms) exchange between different conformations of A_2A_R.

The A_2A_R conformational dynamics was quantitatively analyzed using PET-FCS spectroscopy, by taking advantage of dynamic quenching of a dye attached to residue 229 on TM6 by proximal tyrosines on TM3, TM5, and TM7. PET-FCS analysis revealed fast (sub)microsecond scale protein motions with lifetimes of 150-350 ns, 50-60 μs, and 200-300 μs, respectively. This previously unresolved kinetics is modulated by the binding of orthosteric ligands and of a G-protein mimetic. Surprisingly, the precoupled state promoted by the full agonist shows increased flexibility. One could imagine a wide range of substates that the receptor can sample around this precoupled state, and we believe that this dynamics may facilitate the binding step with the G protein.

The dynamics resolved by FCS is not associated with exchange between active and inactive states of the receptor, but rather describes local fluctuations at the intracellular side of TM3, TM5, TM6 and TM7 in different ligand-biased conformations. These fast intra-state conformational fluctuations of the receptor may represent movement or rotation of helical domains in A_2A_R to facilitate the binding of the G protein or other intracellular effectors. The mini-G protein used here mimics the GTPase domain of the G protein and is specifically engineered to couple to the fully active receptor ^47^. While this mimetic has no nucleotide dependence, the conformational flexibility of A_2A_R revealed by single-molecule fluorescence may be important for facilitating nucleotide exchange (*i*.*e*., GDP to GTP) in the receptor-coupled G protein. Understanding the rich multiscale dynamics of GPCRs will be key to develop a metric that distinguishes the effects of drugs, and their ramifications on the activation processes of G protein and their downstream signaling cascades.

## MATERIALS & METHODS

### Plasmid construction and transformants screening of A_2A_R

The constructs of A_2A_R V229C for fluorescent experiments were same as previously published ^4,5^. The construct, A_2A_R(G119C Q226C), was generated using QuikChange Site-Directed Mutagenesis Kits (Agilent Technologies). The mutant gene was verified by full-length DNA sequencing for target genes (The Center for Applied Genomics, Sick Kids Hospital, Toronto, Canada) with the AOX1 primer pair of PF_AOX1_ and PR_AOX1_ prior to expression transformant screening. Freshly prepared competent cells of a strain of *Pichia pastoris* SMD 1163 (*Δhis4 Δpep4 Δprb1*, Invitrogen) were electro-transformed with individually *PmeI*-HF (New England Biolabs) linearized plasmid using a Gene Pulse II (Bio-Rad). Clone selection was performed as previously described by an in-house two-step transformant screening strategy ^48^ to maximize the expression of receptor for subsequent fluorescence labeling ^49^.

### A_2A_R expression, purification, and labeling

A_2A_R was expressed in yeast according to a previously published protocol ^48^. Briefly, 60 hours after inoculation, yeast cells were harvested and homogenized, and membrane fragments were solubilized in a detergent solution containing 1% MNG-3 (lauryl maltose neopentyl glycol) and 0.02% CHS (cholesteryl hemisuccinate). Solubilized membranes were incubated with Talon resin (Clontech) and then with 100 μM TCEP reducing agent to obtain A_2A_R-bound Talon resin suspended in 50 mM HEPES, pH 7.4, 100 mM NaCl, 0.1% MNG-3, and 0.02% CHS ^50^.

Apo A_2A_R was eluted from the Talon resin with 50 mM HEPES, pH 7.4, 100 mM NaCl, 0.1% MNG-3, 0.02% CHS, and 250 mM imidazole. Sodium chloride and imidazole in the sample were removed by dialysis against 50 mM HEPES, pH 7.4, 0.1% MNG-3, 0.02% CHS for 3 h. A_2A_R was then incubated with BODIPY-FL-iodoacetamide (ThermoFisher Scientific, Cat. no. D6003) for 2 hours for labelling at residue V229C. The XAC-agarose gel, i.e., A_2A_R antagonist xanthine amine congener (XAC) conjugated to Affi-Gel 10 resin, and A_2A_R were subsequently incubated together for 2 hours with gentle nutation. Non-functional A_2A_R and free unreacted BODIPY-FL-iodoacetamide was washed off with 50 mM HEPES, pH 7.4, 100 mM NaCl, 0.1% MNG-3, and 0.02% CHS.

Functional A_2A_R labelled with BODIPY-FL was eluted with 50 mM HEPES, pH 7.4, 0.1% MNG-3, 0.02% CHS, 100 mM NaCl, and 20 mM theophylline. Talon resin was added to the eluted sample and incubated for another 2 h to bind functional A_2A_R-BODIPY-FL, and then was washed extensively with 50 mM HEPES, pH 7.4, 100 mM NaCl, 0.1% MNG-3, 0.02% CHS, and 20 mM imidazole, to remove the theophylline. The functional apo A_2A_R was eluted with 50 mM HEPES, pH 7.4, 100 mM NaCl, 0.1% MNG-3, 0.02% CHS, and 250 mM imidazole, and the sample was dialyzed to remove imidazole for fluorescence experiments. A similar expression, purification, and dye labelling procedure as above was applied for A_2A_R(G119C Q226C), which was designed for smFRET.

### Expression and Purification of R373C-ΔCys-mini-G_s_

The mini-G_s_ construct used in this study is based on the miniG_s_-393 construct described previously ^43^. The miniG_s_-393 construct was modified to remove four cysteine residues (C237S^G.H2.10^, C359I^G.S6.01^, C365V^G.s6h5.02^, and C379V^G.H5.11^) producing a cysteine-less variant. This cysteine-less variant was developed to serve as a background for the introduction of cysteine point mutations for covalent chemical modification. One cysteine residue was introduced on helix 5 in the Ras domain at residue R373^G.H5.05^ to create the R373C-ΔCys-Mini-G_s_ construct used in the current study. Further details regarding the expression and purification can be found in the Supporting Information section 2.

### Single-Molecule FRET

smFRET measurements were performed on a custom-built multiparameter fluorescence microscope with alternating-laser excitation (ALEX) ^35-36^. The donor dye (AF488) was excited with a 473 nm diode-pumped solid-state laser (Cobolt Blue, Market Tech). The acceptor dye (AF647) was excited using a continuous wave 635-nm laser (World StarTech, TECRL-25GC-635-TTL-A). An acousto*-*optic modulator (Isomet, 1205C-2) switches between blue and red excitation with 50 μs duty cycle via a function generator (Tektronix, afg3022b). A multi-channel single-photon counting module (PicoHarp300, PicoQuant, Germany) collects digitally converted photon data from avalanche photodiodes. Green (donor) emission was separated from red (acceptor) emission using a dichroic mirror (FF640-Di01, Semrock) and band-pass filters (BP520/66m and HQ685/80, Chroma), and broadband polarizing cubes further separate the two signals into two orthogonal polarization components.

The smFRET measurements were performed using a concentration of ∼50 pM of sample in 20 mM HEPES (100 mM NaCl, pH 7.4) in the presence of 20 mM cysteamine and 0.01% Tween20 and using blue and red laser excitation with an intensity of ∼25 kW/cm^2^ and ∼12 kW/cm^2^, respectively, at the objective. A custom MATLAB script based on the ‘MLT’ algorithm ^51^ was used to identify fluorescence bursts and sort them into donor-only, acceptor-only and dual-labelled (FRET) populations ^35^. The energy transfer efficiency for each burst, *E*, was estimated from the number of photons detected in the donor, *I*_*D*_, and the acceptor, *I*_*A*_, channels as:

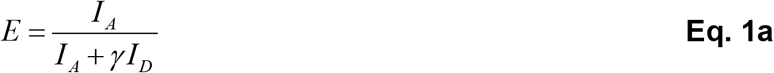

Here the gamma factor, γ, corrects for the difference in detection efficiencies between the donor and acceptor channels (η_Dem_ and η_Aem_), and in the quantum yields of the two dyes (F_D_ and F_A_) ^52^:

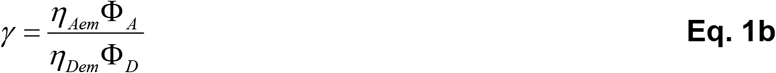

The gamma factor was estimated to be γ = 0.56 ± 0.02 using standard FRET dsDNA samples (see Supporting Information section 1), as described previously ^53^.

The donor-acceptor stoichiometry, S, for the data was calculated as follows,

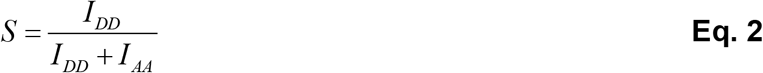

where, *I*_*DD*_, and *I*_*AA*_, represent the spectrally corrected intensities of the donor after donor excitation and of the acceptor after acceptor excitation, respectively. ^53^ Bursts with stoichiometry between 0.2 and 0.8 were used to build the smFRET histograms presented in this work.

Background and spectral-cross-talk corrections were applied using donor-only and acceptor-only bursts, as described previously ^52, 54^. The smFRET histograms were fitted to either one Gaussian distribution or to a sum of two Gaussian distributions. The choice of fitting models was informed by the minimization of reduced-χ^2^ and by the Akaike information criterion (AIC) ^55^. The error margins were estimated using statistical bootstrapping^56^. Briefly, this was achieved via random sampling of *n* bursts from the original experimental pool containing, *N*, smFRET bursts, with replacement (*i*.*e*., some bursts are chosen multiple times, others not at all). As such, synthetic smFRET histograms are generated from each subset containing *n* bursts and each were fit to one or two Gaussians. This algorithm was repeated 2000 times. The standard deviation of the array of best fit parameters from the 2000 synthetic histograms were then taken as the error margin for reported parameter.

### Fluorescence lifetime and anisotropy decay

Fluorescence lifetime and anisotropy decay measurements were performed using the custom-built multiparameter fluorescence microscope (see above) and the data was fit using custom MATLAB code ^36^. BODIPY-FL was excited with 480-nm femtosecond pulses obtained by frequency-doubling the output of a tunable Ti:Sapphire laser (Tsunami HP, Spectra Physics, USA). Time-and polarization-resolved curves were obtained by binning the time delay between excitation and detection events for photons with distinct polarizations (parallel and perpendicular) compared to the polarization of the excitation laser. Measurements were performed with a concentration of ∼100 nM receptor in 20 mM HEPES (100 mM NaCl, pH 7.4) for the ligand-free (apo) receptor, and in the presence of saturating amounts (100 μM) of ligands. All samples were excited at an intensity of 0.1 kW/cm^2^.

Fluorescence polarization anisotropy is defined using the parallel, *I*_*¦*_, and perpendicular *I*_*-*_, components of the detected fluorescence emission as:

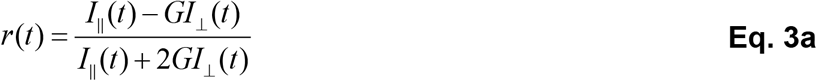

where *G* is a factor that corrects for different sensitivities in detecting signals with different polarizations ^57^. Anisotropy decay time constants of the free and receptor-attached BODIPY-FL, *i*.*e*., rotational correlation times, were estimated by fitting the experimental anisotropy curve *r (t)* to a multi-exponential function:

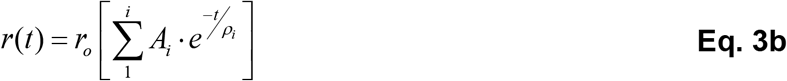

where *r*_o_ is the fundamental anisotropy of the fluorescence particle (∼0.4) and *A*_i_ is the fraction of species with rotational correlation time *ρ*_*i*_. ^57^

The fluorescence lifetimes of the free and the receptor-attached BODIPY-FL were estimated by fitting the isotropic decay curve, i.e., the denominator in **Eq. 3a**, to a multi-exponential decay function:

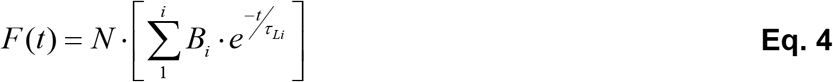

where *N* is the total number of photon counts detected and *B*_*i*_ is the fraction of species with the lifetime *τ*_*Li*_ ^36^. The quality of the fitting and the model validation were evaluated by minimization of χ^2^ and the Akaike information criterion (AIC) ^55^. The standard deviation of the fitting parameters was estimated by parametric bootstrapping ^56^.

### Fluorescence Correlation Spectroscopy

FCS measurements were performed on A_2A_R-V229C-BODIPY-FL using a custom-built fluorescence confocal microscope, described in detail elsewhere ^58^. For each experiment, a solution of 30 μL containing 10 nM receptor in 20 mM HEPES (100 mM NaCl, pH 7.4 supplemented with 20 mM cysteamine and 2 mM Trolox) was cast on a plasma-cleaned glass coverslip (VMR, cat. No. CA48366-249-1) and excited at 488 nm at ∼0.2 kW/cm^2^ using a blue laser (TECBL-488nm; WorldStarTech). FCS measurements were repeated under the same excitation/detection conditions for the ligand-free (apo) receptor, and in the presence of saturating amounts (100 μM) of an inverse agonist (ZM241385), a partial agonist (LUF5834), and a full agonist (NECA) w/wo a G-protein mimetic (mini-G_s_). The resulting pseudo-autocorrelation curves were fitted to the following mathematical model ^33^:

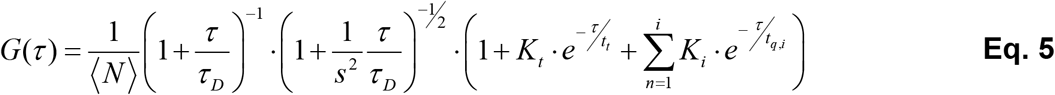

where ⟨*N*⟩ is the average number of molecules in the detection volume (which is characterized by the width *ω*_o_ ellipticity *s*, both measured using samples with known diffusion properties ^58^), τ_D_ is the average diffusion time, *t*_t_ and *K*_t_ are the triplet-state lifetime and amplitude, respectively, and *tt*_q,i_ and *K*_i_ are quenching lifetimes and corresponding amplitudes, respectively.

The hydrodynamic radius (*R*_*H*_) of the fluorescent molecules is estimated by using the FCS-derived diffusion time τ_D_ using the Stokes-Einstein equation:

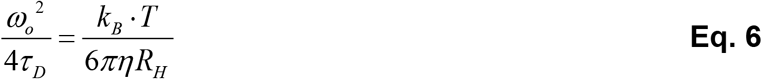

where *ω*_o_ is the width of the confocal detection volume, *k*_*B*_ is the Boltzmann constant, *η* is the viscosity of the solution, and *T* is the temperature. All FCS curves under different ligand-conditions were globally using custom MATLAB code, where the triplet lifetime of the BODIPY-FL was a shared parameter. The goodness of the fit and the number of free parameters in FCS fitting models were assessed using the minimization of residuals (χ^2^) and the AIC criterion ^55^. For clarity, the diffusion component of the FCS curve was subtracted from the fitted curve and the raw FCS data, leaving only the triplet and quenching components, as shown in **Figure 3**.

### BODIPY-FL simulations

The initial structure for an 11-residue segment of A_2A_R (T224 to S234) was taken from PDB: 2YDO.^14^ The V229C mutation and the conjugated dye were built with the molefacture plugin to VMD.^59^ Parameters were constructed with the LigParGen web server ^60^, supplemented with backbone parameters from cysteine and the C-C-S-C dihedral parameter from methionine. The dye’s boron atom was replaced with a carbon atom to circumvent limitations of the LigParGen web server. No attempt was made to enforce boron-like geometry, though inaccuracies stemming from this simplification are likely minor due to regional rigidity. Three 2.5 μs simulations of this system were conducted in a cubic box with ∼6,000 water molecules and 100 mM excess KCl. To maintain helical structure throughout these simulations, residue restraints with force constants of 10^3^ kJ/mol/nm^2^ were applied to all C_α_ atoms. To quantify the spatial separation of the dye from its protein attachment, distances, *r*, were measured from the modified cysteine C_β_ atom to the center of geometry of non-hydrogen atoms in the dye’s aromatic group (C1, C2, C4, C5, N1, C7, C8, N2, and B1, where B1 represents the boron that we modeled as carbon). Further details regarding these simulations can be found in the Supporting Information Section 3.

### Molecular Dynamics Simulations of A_2A_R

We conducted six MD simulations for each of seven different initial receptor configurations, which were obtained from crystal structures containing either the inverse agonist ZMA (PDBs: 3PWH and 4EIY), ^61-62^ the endogenous agonist adenosine (PDB: 2YDO), ^14^ the synthetic agonist UK-432097 (PDB: 3QAK), ^13^ or the agonist N-ethyl-5’-carboxamido adenosine (NECA) and an engineered G protein (PDBs: 5G53 and 6GDG). ^47,63^We used both chains A and B from the 5G53 structure independently. None of our simulations included ligand, G protein, or receptor modifications employed for crystallization. The simulated sequence was S6 to V307 (i.e., ΔM1-G5 and ΔL308-S412). Residue identifiers always corresponded to the receptor’s full-length wild-type sequence (UNIPROT ID: P29274). The receptor was not glycosylated, all titratable residues were in their standard states for pH 7, and the protein backbone termini were zwitterionic. Further details regarding the MD simulations can be found in the Supporting Information Section 3.

## ASSOCIATED CONTENT

### Supporting Information

Descriptions of smFRET control samples, expression, and purification of mini-G, and molecular dynamics simulations. Additional tables and figures for MD data, FAD, and FCS control experiments.

## AUTHOR INFORMATION

### Competing Financial Interest

The authors report no competing financial interest.

## ACKNOWLEDGEMENTS

This work was supported by the Natural Sciences and Engineering Research Council of Canada (Grant No. RGPIN 2017-06030 to C.C.G.) and by Canadian Institutes for Health Research (Grant No. MOP-43998 to R.S.P.) C.N. was funded by U.S. DOE Laboratory Directed Research and Development (LDRD) funds. Computations used resources provided by the Los Alamos National Lab Institutional Computing Program, which is supported by the U.S. DOE National Nuclear Security Administration under Contract No. DE-AC52-06NA25396.

## Notes

### Competing Interest Statement

The authors have declared no competing interest.

