## Supporting Information for "Ligand Modulation of the Conformational Dynamics of the A_2A_ Adenosine Receptor Revealed by Single-Molecule Fluorescence"

GPCRs, single-molecule fluorescence, smFRET, FCS, photoinduced electron transfer, molecular dynamics, conformational dynamics

**SUPPORTING INFORMATION SECTION 1:**

### Single-Molecule Förster Resonance Energy (smFRET)

#### 1.1 Design Rationale for T119-Q226 $A_{2A}R$ Mutant for smFRET:

To track inward and outward movements from TM6, several published crystal structures of the $A_{2A}R$ were evaluated to find the positions which provided most dynamic changes for smFRET. Overlapping the crystal structures of the inverse agonist-, full agonist-, and mini-G-bound crystal structures of $A_{2A}R$ (*i.e.*, 3EML, 2YDO, and 5G53, respectively) revealed that position Q226C on TM6 showed a pronounced displacement and rotated outwards whereas position T119C located on the alpha helix in intracellular domain II (ICL2) near TM4, remained relatively immobile (**Figure S1**). The displacement of the between these two points corresponded to ~10 Å (**Figure S1**), which is similar with the results suggested by our smFRET measurements.

#### 1.2 Double-stranded DNA: smFRET Corrections and Static Limit Controls

To obtain the $\gamma$ correction factor for smFRET measurements (**Eq. 1a-b**, main text, Materials & Methods), we designed two 30 base pair (bp) double-stranded DNA (dsDNA) with expected FRET efficiencies at ~30% and ~80%, respectively. Complementary single-stranded (ss) DNA strands were synthesized (Integrated DNA Technologies (IDT), Inc., Coralville, Iowa, USA), with one strand containing the donor (fluorescein) via a thymine ( **T** ) ring on position five, and the other strand containing the acceptor (AF647) attached to the 5'-end . Donor- and acceptor-labelled oligonucleotides were annealed to obtain dsDNA with desired donor-acceptor base-pair (bp) separations. The annealing buffer contained 10 mM Tris pH 7.5, 50 mM NaCl and 1 mM EDTA. Equimolar amounts (100 μM) of the two strands were heated to 95 $^{o}$C for 2 minutes using a thermocycler (Biometra TPersonal Thermocycler), followed by slow cooling to 25 $^{o}$C for 45 minutes, after which the sample was aliquoted and frozen at -20 $^{o}$C.

The smFRET control measurements were performed on two dsDNA samples with separations of 10bp and 17bp between the donor (fluorescein) and the acceptor (AF647) fluorophores, which were attached via a six-, and a five-carbon linker, respectively. The experiments were conducted using the same excitation power as was used for $A_{2A}R$ (donor and acceptor lasers intensities of ~25 kW/cm$^2$ and ~12 kW/cm$^2$, respectively), and in a buffer solution (50 mM TRIS, 1 mM EDTA, pH 8.5, 150 mM NaCl) supplemented with 10 mM cysteamine. Single-peak FRET histograms were observed for both 10 bp and 17 bp dsDNA samples, having mean efficiencies $\langle \boldsymbol{E} \rangle$ of 80 ± 1 % and 29 ± 1 %, and widths **$\Delta E_{FWHM}$** of 19 ± 1 % and 38 ± 1 %, respectively. In addition to γ correction for smFRET, the width of the FRET histogram for the 10 bp dsDNA (pink band in **Figure 1b**) serves as a quasi-static limit for the exchange broadening in $A_{2A}R$, i.e. the minimum FRET width in the absence of structural dynamics of the labelled biomolecule.

An upper limit for the smFRET width due to shot noise, **$\Delta E_{shot\text{-}noise}$**, can be computed using the expression:

$$\Delta E_{shot-noise} = 2 \cdot \frac{\sqrt{E \cdot (1-E)}}{\sqrt{N}} \qquad \textbf{Eq. S1}$$

where $E$ is the FRET efficiency and $N$ is the average number of photons per burst. Four values were calculated using mean efficiencies $\langle \boldsymbol{E} \rangle$ and photon numbers for the $A_{2A}R$ samples. The blue band in **Figure 1b** represents a spline (polynomial) interpolation of these points, with the width obtained by propagating errors in **Eq. S1.**

#### 1.3 Förster Radius in $A_{2A}R$: Limitations of the Donor-Acceptor Orientation Factor $\kappa^2$

The efficiency of the Förster resonance energy transfer $E$ between a donor and an acceptor fluorophore depends on the distance $R$ between the fluorophores according to the expression:

$$E = \frac{1}{1+\left(R/R_0\right)^6} \qquad \textbf{Eq. S2}$$

The Förster radius $R_o$ is calculated according to the expression:

$$R^6_o = \frac{9000 \cdot \ln(10) \cdot \Phi_D \kappa^2}{128\pi^5 N n^4} \int_0^\infty f_D(\lambda)\varepsilon_A(\lambda)\lambda^4 d\lambda \qquad \textbf{Eq. S3}$$

where $J$ is the normalized spectral overlap integral $J = \int_0^\infty f_D(\lambda)\varepsilon_A(\lambda)\lambda^4 d\lambda$, $\Phi_D$ is the fluorescence quantum yield of the donor , $N$ is Avogadro's number , $\kappa^2$ is the orientation factor, $n$ is the refractive index, $\varepsilon_A$ is the extinction coefficient of the acceptor, and $\lambda$ is the wavelength. For the AF488 and AF647 dyes conjugated to $A_{2A}R$, absorption and emission spectra were measured and used to calculate the spectral overlap integral using open source software (UV-Vis-IR Spectral Software 1.2, FluorTools). We estimated the quantum yield of AF488 as $\Phi_D = 0.72 \pm 0.02$ using the comparative method [1].

To assess whether the isotropic value $\kappa^2 = 2/3$ is a good approximation for the $A_{2A}R$ samples, we measured the steady-state anisotropy or donor- and acceptor-labelled receptors. The values obtained from averaging over hundreds of single-molecule bursts were $r_{AF488} = 0.34 \pm 0.02$ for the donor and $r_{AF647} = 0.024 \pm 0.005$ for the acceptor [2]. Using these values, we calculated the lower and the upper limit of the orientation factor, $\kappa_{min}^2$ and $\kappa_{max}^2$, respectively[3]:

$$\kappa^2_{\min} = \left[ 2/3 \right] \cdot \left( 1 - \frac{5}{2} \left[ \frac{\langle r_{AF\,488} \rangle + \langle r_{AF\,647} \rangle}{2} \right] \right) = 0.363 \quad \textbf{Eq. S4}$$

$$\kappa^2_{\max} = \left[ 2/3 \right] \cdot \left( 1 + \frac{5}{2} \left[ \langle r_{AF\,488} \rangle + \langle r_{AF\,647} \rangle \right] \right) = 1.27 \quad \textbf{Eq. S5}$$

These limits correspond to a minimum and maximum $R_o$ of 45 and 56 Å, respectively. This range exceeds the typical error margin for $R_o$, which depends on the error in measuring the quantum yield of the donor, $\Phi_D$, and in estimating the donor-acceptor spectral overlap integral.

### SUPPORTING INFORMATION SECTION 2:

### Expression, Purification, and Labelling of $A_{2A}R$ and Mini-$G_s$

#### 2.1 $A_{2A}R$ expression, Purification and Labeling.

A single yeast colony on YPD plates with a desirable expression level was inoculated into 4 mL of YPD medium and cultured overnight. The medium was then inoculated into 200 mL of BMGY medium (1% (w/v) yeast extract, 2% (w/v) peptone, 1.34% (w/v) Yeast nitrogen base (YNB) without amino acids, 0.00004% (w/v) biotin, 1% (w/v) glycerol, 0.1 M PB (phosphate buffer) at pH 6.5) and culture for an additional 24 h. Cells were spun down at 4000 rpm for 10 min and resuspended in 1 L of BMMY medium (1% (w/v) yeast extract, 2% (w/v) peptone, 1.34% (w/v) YNB without amino acids, 0.00004% (w/v) biotin, 0.5% (w/v) methanol, 0.1 M phosphate buffer at pH 6.5, 0.04% (w/v) histidine and 3% (v/v) DMSO, 10 mM theophylline) at 20 °C. Methanol (0.5% (v/v)) was added every 12 h for inducing the receptor expression.

Cell pellets were harvested by centrifugation at 4000 rpm for 10 min after 60 hrs induction and washed with 50 mM HEPES, pH 7.4 before the addition of breaking buffer (50 mM HEPES, pH 7.4, 100 mM NaCl, 2 mM EDTA, 10% glycerol, 100 μM theophylline). The cell pellets were applied to heavy duty vortex for 2 hrs at 4°C to release the membrane fractions. Undisrupted cells (debris) were separated from the membrane suspension by centrifugation (8,000 g) for 30 min. The supernatant was collected and further centrifuged at 100,000 xg for 1 h. The precipitated cell membrane was then immediately dissolved in 50 mM HEPES, pH 7.4, 100 mM NaCl, 1% MNG-3 (lauryl maltose neopentyl glycol) and 0.02% CHS (cholesteryl hemisuccinate), 100 μM theophylline, and 20 mM imidazole under continuous agitation for 1–2 hrs at 4°C till the solution was transparent.

Subsequently, Talon resin (Clontech) was added to the solubilized membranes and incubated for 2 h. The $A_{2A}$R-bound Talon resin was washed with 50 mM HEPES buffer, pH 7.4, containing 100 mM NaCl, 0.1% MNG-3, and 0.02% CHS and resuspended in the same buffer, followed by addition of 100 μM TCEP reducing agent and incubation additional 20 min. TCEP was washed out immediately with a buffer consisting of 50 mM HEPES, pH 7.4, 100 mM NaCl, 0.1% MNG-3, and 0.02% CHS. The $A_{2A}$R-bound Talon resin was then incubated with BODIPY-FL-iodoacetamide (ThermoFisher Scientific, Cat. no. D6003) for overnight labeling. The labeled $A_{2A}$R was packed onto a disposal column and washed 2 columns buffers of 50 mM HEPES, pH 7.4, 100 mM NaCl, 0.1% MNG-3, and 0.02% CHS. The $A_{2A}$R was then eluted from Talon resin column using 50 mM HEPES, pH 7.4, 100 mM NaCl, 0.1% MNG-3, and 0.02% CHS, 300 mM imidazole.

The eluted fractions were concentrated and incubated with XAC-agarose gel (antagonist xanthine amine congener (XAC) conjugated to Affi-Gel 10 resin) for 2 hrs. Non-functional $A_{2A}$R and residual free unreacted BODIPY-FL- iodoacetamide was washed off with 50 mM HEPES, pH 7.4, 100 mM NaCl, 0.1% MNG-3, and 0.02% CHS. Functional $A_{2A}$R was eluted with 50 mM HEPES, pH 7.4, 0.1% MNG-3, 0.02% CHS, 100 mM NaCl, and 20 mM theophylline. Talon resin was added to the eluted sample and incubated for another 2 hrs to bind functional $A_{2A}$R. Conjugated $A_{2A}$R was washed extensively with 50 mM HEPES, pH 7.4, 100 mM NaCl,

0.1% MNG-3, 0.02% CHS, and 20 mM imidazole, to remove all theophylline. The functional apo $A_{2A}R$ was eluted with 50 mM HEPES, pH 7.4, 100 mM NaCl, 0.1% MNG-3, 0.02% CHS, and 250 mM imidazole, and the sample was dialyzed against 50 mM HEPES, pH 7.4, 100 mM NaCl, 0.1% MNG-3, 0.02% CHS to remove imidazole for fluorescent experiments. A similar procedure as above was applied for the construct of $A_{2A}R$_T119C_Q226C which was designed for smFRET in this study.

#### 2.2 Expression and Purification of R373C-ΔCys-mini-$G_s$

R373C-ΔCys-mini$G_s$ was expressed and purified according to the methodology described by Carpenter & Tate[4], with some modifications. Plasmid DNA with the mini-$G_s$ construct was transformed into *E. coli* strain BL21(DE3) cells made chemically competent. Cells were cultured in LB media in 2.8 L Fernbach baffled flasks at 37°C shaking at 200 rpm until an $OD_{600}$ of 0.7-0.8 was reached. Protein expression was induced with the addition of 50 μM IPTG and the temperature reduced to 22°C. Cells were harvested 20 h post-induction by centrifugation at 5000g for 5 min, flash-frozen in liquid nitrogen and stored at -20°C. Cell pellets were resuspended in binding buffer (50 mM HEPES, pH 7.4, 300 mM NaCl, 10% glycerol, 2 mM $MgCl_2$, 5 mM imidazole, 50 μM GDP) and supplemented with 1 mM PMSF, 5 mM 6-aminocaproic acid, 5 mM benzamidine hydrochloride, DNAse I (kk units), Lysozyme (kk units), and 10 mM β-mercaptoethanol. Resuspended cell pellets were lysed on ice by sonication on a kk at 25 dB with 10 s on/off intervals for 2 min and clarified by centrifugation at 12000g for 30 min.

The clarified lysate was passed through a gravity flow column loaded with 3 mL of cOmplete His-Tag Purification resin multiple times. The column was washed with 5 column volumes of wash buffer (50 mM HEPES, pH 7.4, 300 mM NaCl, 10% glycerol, 2 mM $MgCl_2$, 10 mM imidazole, 50 μM GDP, 10 mM β-mercaptoethanol) followed by elution with 3 column volumes of elution buffer (50 mM HEPES, pH 7.4, 300 mM NaCl, 10% glycerol, 2 mM $MgCl_2$, 250 mM imidazole, 50 μM GDP, 10 mM β-mercaptoethanol). The eluted fraction was spin concentrated to 5 mL with a 10 kDa MWCO Amicon centrifugal filter and was then buffer exchanged four times with dialysis buffer (50 mM HEPES, pH 7.4, 300 mM NaCl, 10% glycerol, 2 mM $MgCl_2$, 50 μM GDP, 1 mM TCEP) and concentrated to ~4 mL using the same filter. The N-terminal his-tag was not cleaved during the current study. The concentrated protein sample was applied to a HiLoad 16/600 Superdex 200 pg column equilibrated with gel filtration buffer (50 mM HEPES, pH 7.4, 100 mM NaCl, 10% glycerol, 2 mM $MgCl_2$, 1 μM GDP, 0.5 mM TCEP). The peak fractions were collected and analyzed using SDS-PAGE gel-electrophoresis (**Figure S9**). The collected from was concentrated to 150 μM, aliquoted and flash-frozen in liquid nitrogen. The frozen aliquots were stored at -80°C.

### SUPPORTING INFORMATION SECTION 3:

### Molecular Dynamics Simulations

#### 3.1 Simulations of $A_{2A}R$ in Different Extracellular Ligand States

To build simulation systems, extracellular ligands, water molecules, and other solvents resolved in crystal structures were removed. Missing wild-type backbone atoms were modeled with the program Loopy. [5,6] Missing side chain atoms and side chain reversions to wild-type were modeled with the program SCWRL4.[7] Each repeat used a different model of missing receptor atoms. Disulfide bonds (C71-C159, C74-C146, C77-C166, and C259-C262) and all hydrogen atoms were placed with the GROMACS tool pdb2gmx.[8] A unique bilayer conformation was constructed for each simulation by extracting a randomly selected snapshot from a 300 ns simulation of a neat POPC bilayer with 166 lipids per leaflet, which was initially constructed with the CHARMM-GUI[9] membrane builder.[10]

The receptor was oriented for insertion using the program LAMBADA,[11] and embedded in the bilayer using 20 cycles of the InflateGRO2 routine[11] with double-precision GROMACS steepest descent energy minimization. During this procedure, 11-17 lipids were removed from each leaflet, allowing the final numbers to be asymmetric. Each system was hydrated with ~85 waters per lipid and 100 mM excess KCl, disallowing placement of water or ions in the bilayer's hydrophobic core. Subsequently, water, lipids, and ions were relaxed over 30 ns by sequential 5-ns simulations using position restraints on receptor heavy atoms with force constants of $10^4$ and $10^3$ kJ/mol/nm$^2$, followed by restraints on receptor $C_\alpha$ atoms with force constants of $10^3$, $10^2$, 10, and 1 kJ/mol/nm$^2$.

All restraints were removed for production simulation. Simulations were conducted with mixed-precision (SPFP[12]) AMBER 16 software.[13] Protein was modeled by the CHARMM36m force field[14] and lipids were modeled by CHARMM36.[15] The water model was TIP3P[16] with CHARMM modifications.[17] AMBER formatted topologies were obtained with the gromber tool of ParmEd from AmberTools 16 after initial topology construction with GROMACS 5.1.2.[18] Water molecules were rigidified with SETTLE[19] and other covalent bond lengths involving hydrogen were constrained with SHAKE[20] (tolerance $=10^{-6}$ nm).

Lennard-Jones (LJ) interactions were evaluated using an atom-based cutoff with forces switched smoothly to zero between 1.0 and 1.2 nm. This is the recommended LJ cutoff for the CHARMM36 protein force field.[21,22] Note that the CHARMM36 lipid force field was parameterized with LJ switching between 0.8 and 1.2 nm.[15] Coulomb interactions were calculated using the smooth particle-mesh Ewald method[23,24] with Fourier grid spacing of 0.08 to 0.10 nm and fourth order interpolation. Temperature and pressure were controlled by velocity Langevin dynamics[25] at 310 K with a coupling constant of 1 ps and semi-isotropic coupling to Monte Carlo barostats[13] at 1.01325 bar with compressibilities of $4.5\times10^{-5}$ bar$^{-1}$, respectively. The integration time step was 4 fs with hydrogen mass repartitioning.[26] Non-bonded neighbor-lists were built to 1.4 nm and updated heuristically.

To estimate the precision of our results, the standard error of the mean is obtained using **Eq.S6**, for $N$ repeat simulations with mean values $\mu_i$ and overall mean $\mu$.

$$\mathrm{SE} = \frac{1}{\sqrt{N}}\sqrt{\sum_{i=1}^{N}\left(\mu_i - \mu\right)^2 \Big/ \left(N-1\right)} \qquad \textbf{Eq. S6}$$

To quantify relaxation, we estimated exponential autocorrelation times, $\tau_{\mathrm{acorr}}$, for values of the distance between the BODIPY modified cysteine $C_\beta$ atom and the center of geometry of non-hydrogen atoms in the dye's aromatic group, $l$. The normalized autocorrelation function was evaluated according to **Eq. S7**, where the mean and standard deviation of the sample of $l$ are represented by $\mu$ and $\sigma$, respectively, $\Delta t$ is the time interval, and angular brackets indicate averaging over all possible initial times $t$ (*i.e.*, $t \leq t_{\mathrm{MAX}} - \Delta t$, for a simulation of $t_{\mathrm{MAX}}$).

$$C\left(\Delta t\right) = \left\langle \frac{\left[l\left(t\right)-\mu\right]\times\left[l\left(t+\Delta t\right)-\mu\right]}{\sigma^2} \right\rangle_t \qquad \textbf{Eq. S7}$$

The autocorrelation function, $C(\Delta t)$, was fit to an exponential decay of the form $\exp\left(-\Delta t/\tau_{\mathrm{ac}}\right)$ to obtain $\tau_{\mathrm{acor}}$.

### SUPPORTING FIGURES

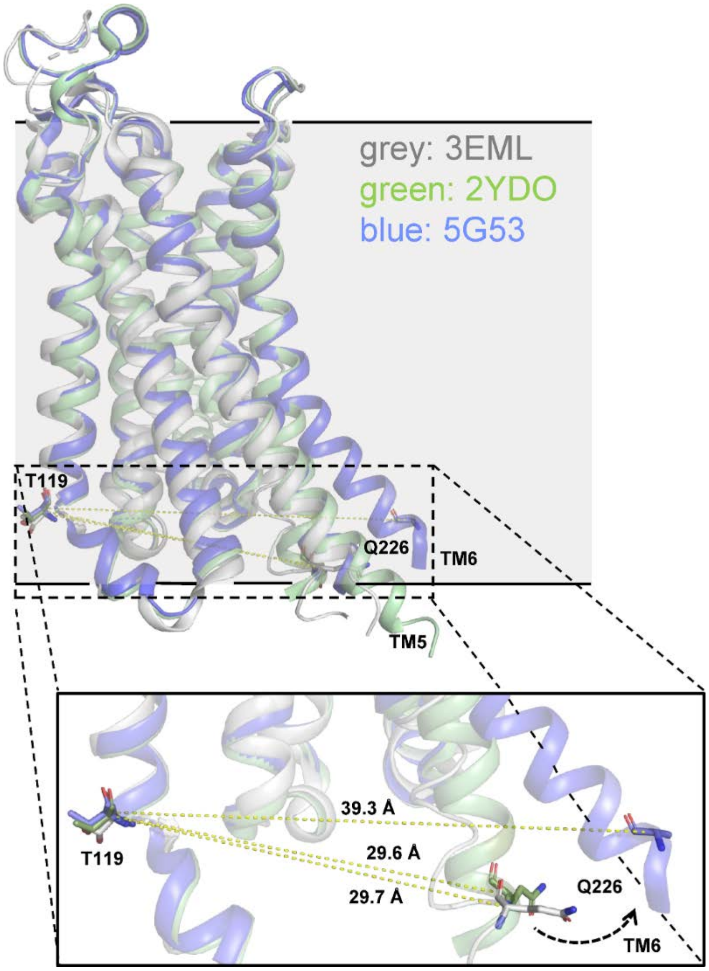


**Figure S1. Rationale for the design of the $A_{2A}R$ mutant for smFRET experiments.** Overlaid crystal structures of $A_{2A}R$ with ZM241385 bound (3EML), NECA bound (2YDO), and when coupled to mini-$G_s$ (5G53). Residue T119 (TM4) remains relatively fixed as shown in the figure, while Q226 (TM6) is mobile. As a result, both of these sites were mutated into cysteine residues to attach maleimide-functionalized donor and acceptor fluorophores (AF488 and AF647) for smFRET measurements.

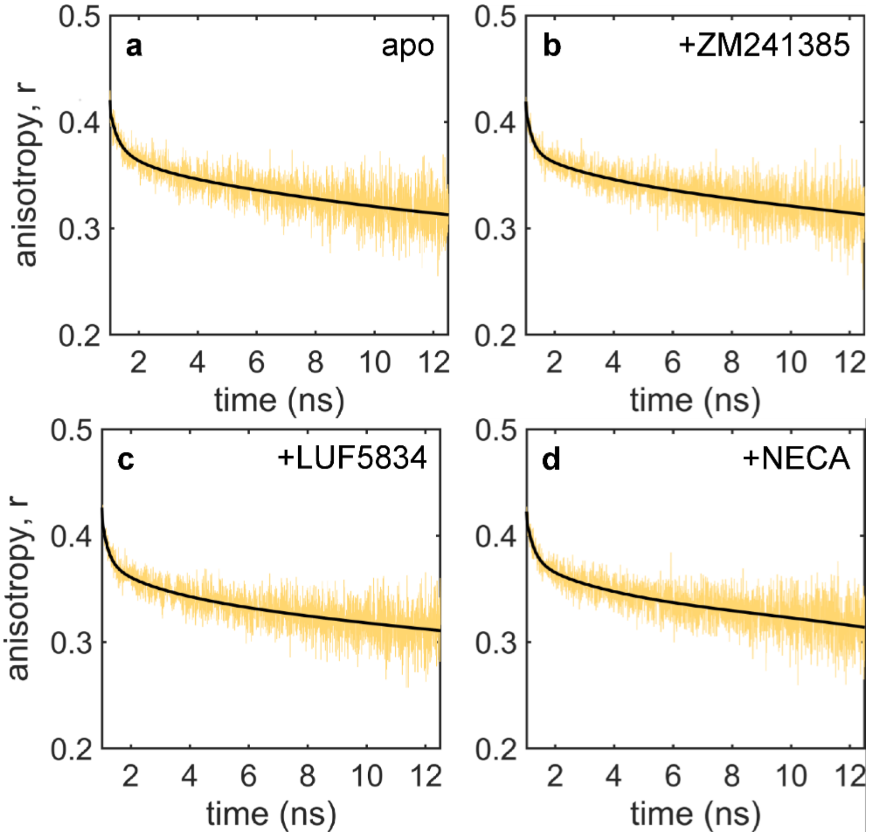


**Figure S2. Time-resolved fluorescence anisotropy measurements of $A_{2A}R$.** Fluorescence anisotropy decay curves of BODIPY-FL-labelled receptors at residue 229, in the apo state **(a)**, and in the presence of **(b)** the inverse agonist ZM241385, **(c)** the partial agonist LUF5834, and **(d)** the full agonist NECA. Solid black lines represent the fitting curves according to **Eq. 3b.** The experiments were performed on 100 nM solutions of labelled receptors w/wo saturating amounts of ligands (100 μM) using pulsed 480-nm laser excitation. The fitting results are summarized in **Table 2.**

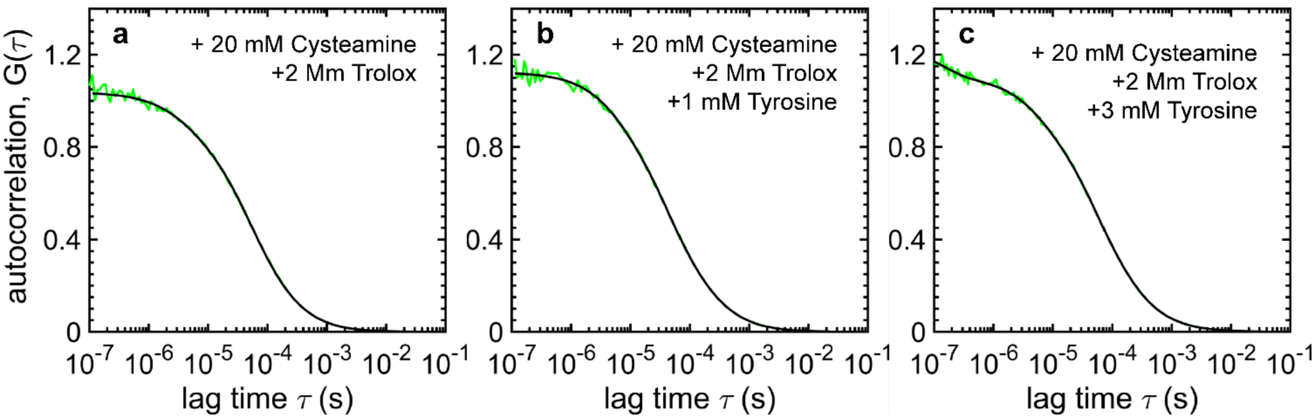


**Figure S3. PET-FCS experiments on free BODIPY-FL in the presence of tyrosine.** FCS data (green) and fitting to **Eq. 5** (black) of free BODIPY-FL **(a),** and of BODIPY-FL in presence of 1 mM **(b)** and 3 mM **(c)** of tyrosine. Experiments were performed on 10 nM dye in a buffer solution (50 mM HEPES, pH 7.4, 100 mM NaCl) with 0.1% MNG-3, 0.02% CHS, and photoprotectant (20 mM cysteamine and 2 mM Trolox). Each sample was excited by a continuous-wave laser at 488 nm and 0.2 kW/cm$^2$. The fitting results are summarized in **Table S1**.

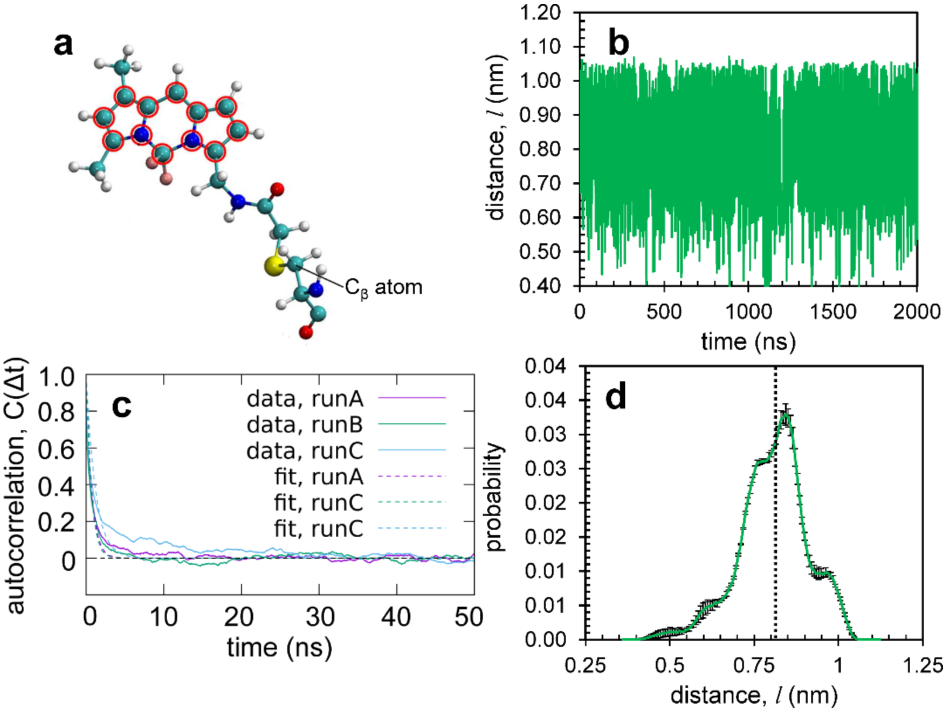


**Figure S4. Molecular dynamics simulations of BODIPY-FL.** Spatial extension of the **(a)** BODIPY-FL molecule (with boron replaced by carbon) attached to position 229 within a small peptide segment of $A_{2A}R$ (224-234). The red circles in **(a)** indicate the 12 atoms that were used to compute the center of geometry of the aromatic system of BODIPY-FL. **(b)** Distance (*l*) between the indicated $C_\beta$ atom in **(a)** and the computed center of geometry of BODIPY-FL, illustrated as a representative time series (green line) from 3 independent 2-μs simulations of. **(c)** For each time series, distance autocorrelation decays, $C(\Delta t)$ conf. **Eq. S7**, were calculated (solid colored lines) and fitted to mono-exponential functions (dashed colored lines); the estimated autocorrelation lifetime, $\tau_{acor}$, describes the timescale of dye linker relaxation. **(d)** Probability distribution of the distance *l* (green line), including the standard deviation from 3 replicates (black bars) and the mean value, 0.81 ± 0.01 nm (vertical dashed line).

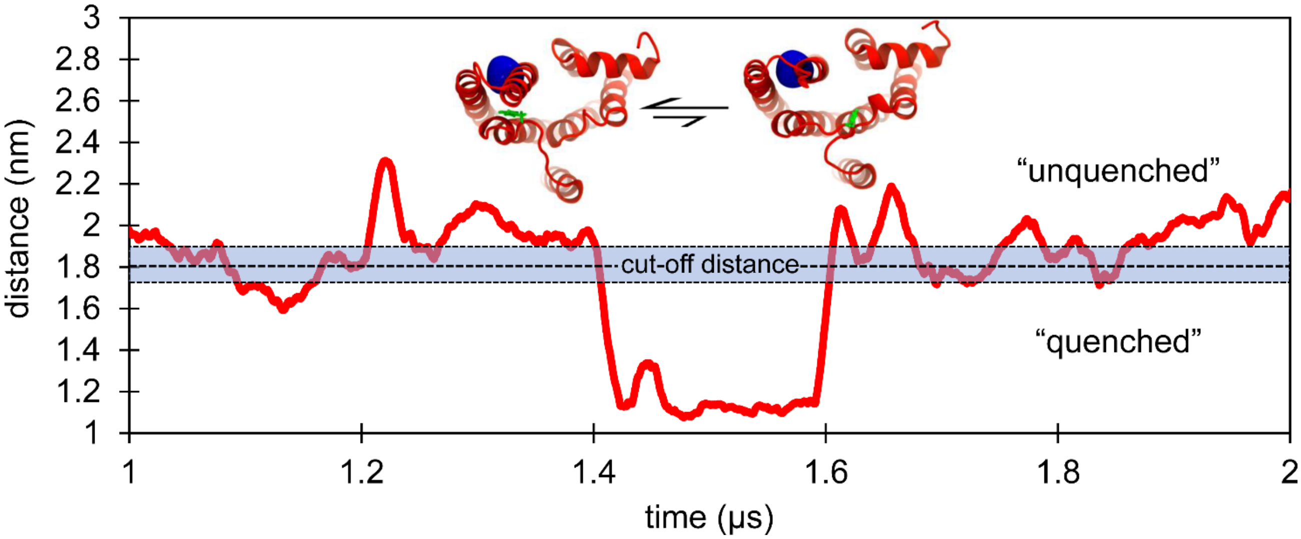


**Figure S5. MD-based estimation of the rate of PET for BODIPY-FL in $A_{2A}R$.** A representative molecular dynamics trajectory of the distance (red line) between the BODIPY-FL anchoring point on residue 229 ($C_\beta$) and the center of mass of an aromatic residue in $A_{2A}R$. The trajectory is overlaid with the PET quenching cut-off distance characteristic of the BODIPY-FL-linker system, $d_q = 1.8 \pm 0.1$ nm (blue shaded region). The rate of quenching/dequenching was estimated by counting the frequency of crossing between the two quenching zones.

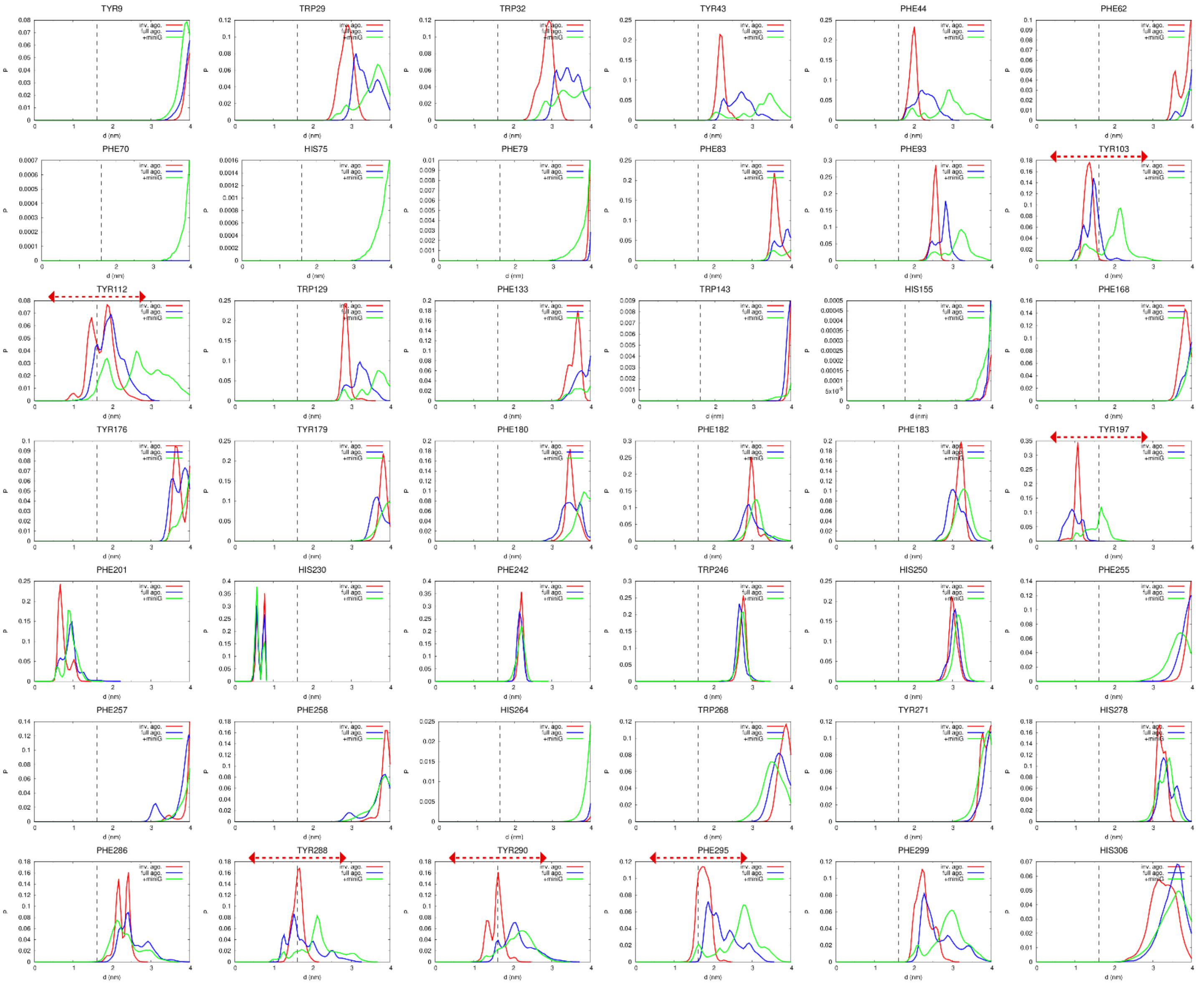


**Figure S6. Identifying aromatic residues in A$_{2A}$R within a feasible distance for PET quenching of BODIPY-FL attached to residue 229.** Probability histograms show the distributions of the distance (*d*) between the V229 C$_{\beta}$ atom to the center of mass of each aromatic side chain. Each subplot represents one aromatic residue. Distributions shown for 12 simulations of structures crystallized with inverse agonist (red), 12 simulations of structures crystallized with full agonist (blue), and 18 simulations of structures crystallized with agonist and mini-G (green). Dashed vertical gray line denotes the cut-off quenching distance $d_q$ = 1.8 nm. Dashed horizontal red line indicates aromatic residues with an appreciably populated distributions on either side of the quenching distance $d_q$.

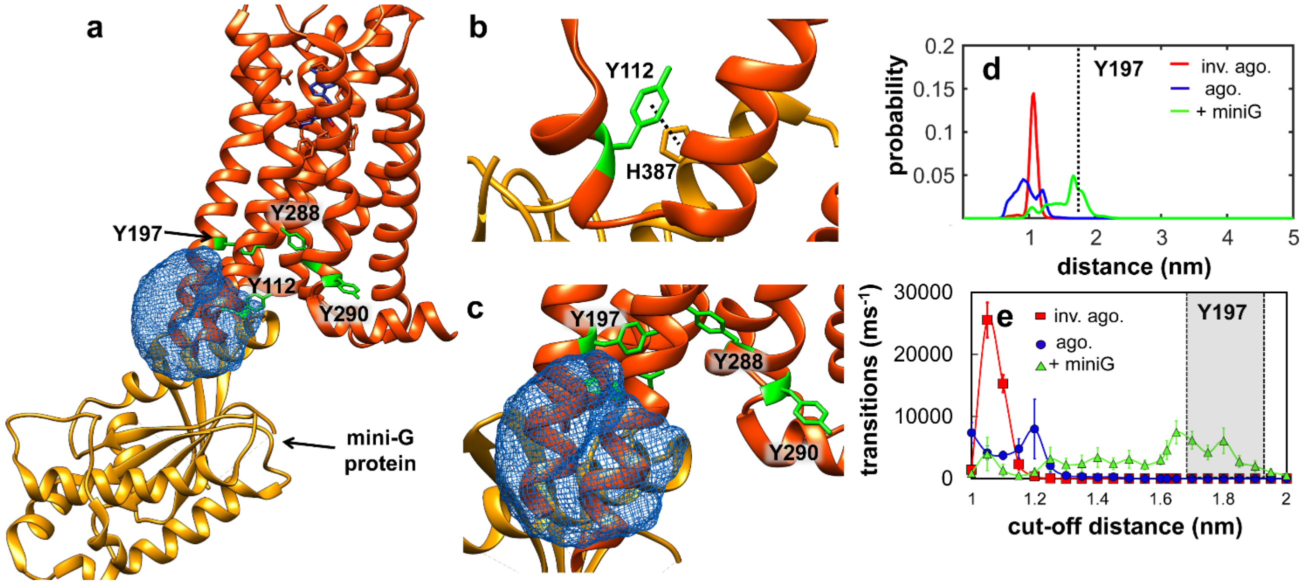


**Figure S7. Molecular dynamics simulations of fluorescently labelled $A_{2A}R$ with mini-G protein. (a)** PDB (5G53) structure used for simulations of receptor with mini-G protein bound, with the computed BODIPY-FL cloud attached from overlaid. **(b)** Zoomed region of interest of **(a)** revealing stabilization of Y112 (green) of the receptor through an interaction with H387 (gold) of mini-G protein. Despite the absence of mini-G in our MD trajectories, simulations initiated from crystal structures of $A_{2A}R$ bound to mini-G frequently retained substantial displacement between TM3 and TM6 and a broken ionic lock (see **Figure S8**). **(c)** Zoomed region of interest in **(a)** containing the tyrosines Y197, Y288 and Y290 (green). Probability distributions for the distance **(d)** between the alpha-carbon of position 229 to the center of mass of Y197 and the frequency of quenching transitions as a function of cut-off distance **(e)**, in presence of the inverse agonist (red), full agonist blue), and full agonist plus mini-G protein (green).

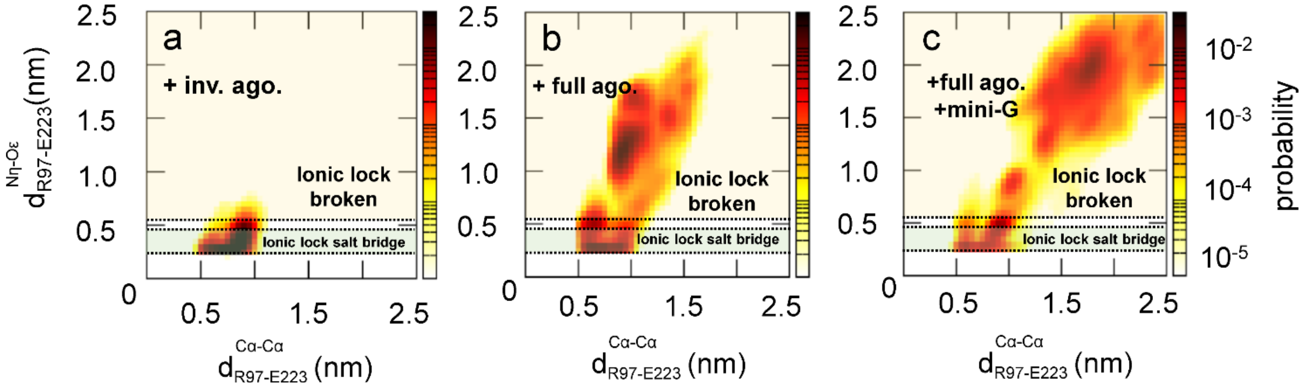


**Figure S8. TM3-TM6 association and R93-E223 ionic lock formation in MD simulations.** 2D probability distributions relating the distance between Cα atoms of TM6 residue E223 and TM3 residue R97, $d^{C\alpha-C\alpha}_{\text{R97-E223}}$ , to the minimum distance between side chain arginine η nitrogen atoms (Nη) and glutamate ε oxygen atoms (Oε) for this residue pair, $d^{N\eta-O\varepsilon}_{\text{R97-E223}}$. The former term, shown on the abscissa, reports on the TM3-TM6 displacement, whereas the latter term, shown on the ordinate, reports on formation of the ionic lock. Data shown reports on ionic lock stability for simulations initiated using crystal structures in which the $A_{2A}R$ was crystalized bound to **(a)** an inverse agonist, **(b)** a full agonist, or **(c)** a full agonist plus mini-G, as shown in **Table S2**. Shaded dashed yellow region **(a-c)** indicate distances were the ionic lock is broken, whereas shaded dashed green regions **(a-c)** indicate the regime for salt bridge formation with the ionic lock intact. Probability scales in **(a-c)** are indicted by the spectrum of colors from white to dark red and follow a logarithmic scale.

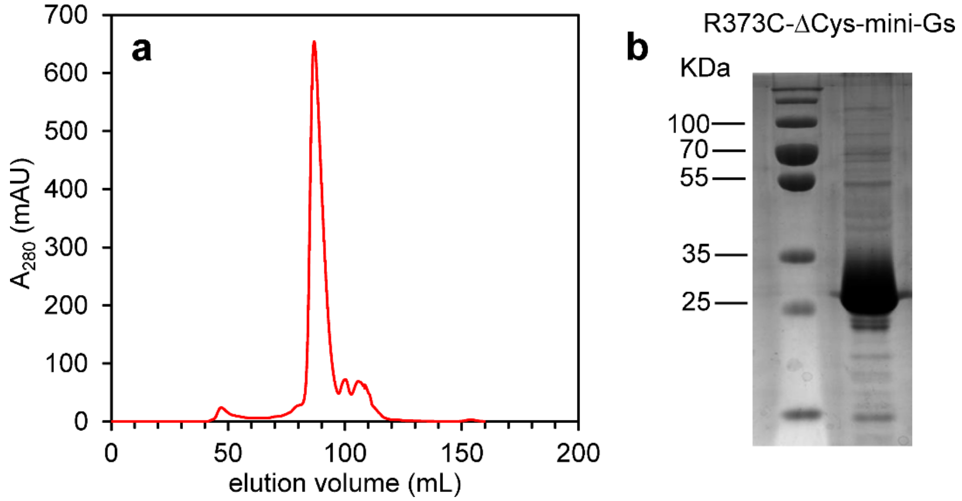


**Figure S9. Purification and purity of mini-G protein. (a)** Size-exclusion chromatogram of elution fractions of purified mini-G protein (R373C-ΔCys, mini-$G_S$), and **(b)** the SDS-PAGE gel electrophoresis image, showing the corresponding molecular weight 28.9 kDa.

### SUPPORTING TABLES

**Table S1.** FCS fitting parameters for free BODIPY-FL in the presence of tyrosine [a,b]

| | BODIPY-FL | + 1 mM Tyrosine | +3 mM Tyrosine |
|---|---|---|---|
| **$t_{trip}$ (μs)** | 3 ± 1 | 6 ± 1 | |
| **$t_{fast}$ (μs)** | — | — | 0.11 ± 0.03 |
| **$t_{int}$ (μs)** | — | 40 ± 2 | 46 ± 2 |
| **$t_{slow}$ (μs)** | — | — | — |

[a] Data was measured on 10 nM solutions of free dye in the presence of 20 mM Cysteamine and 2 mM Trolox
[b] FCS curves were fitted to **Eq. 5**; parametric errors are the standard errors obtained from the fit.

**Table S2.** $A_{2A}R$ structures used for molecular dynamics simulations

| Structure | Chain | Extracellular Ligand [a] | Intracellular Protein [b] | *N* sim. | time (μs) |
|---|---|---|---|---|---|
| **3PWH** | A | inverse agonist | — | 6 | 5 |
| **4EIY** | A | inverse agonist | — | 6 | 5 |
| **2YDO** | A | agonist | — | 6 | 5 |
| **3QAK** | A | agonist | — | 6 | 5 |
| **5G53** | A | agonist | mini $G_s$ | 6 | 5 |
| | B | agonist | mini $G_s$ | 6 | 5 |
| **6GDG** | A | agonist | mini G (trimer) | 6 | 5 |

[a,b] Indicates crystallization conditions; co-crystallized extracellular ligands and intracellular proteins were not included in simulations.
